# Multiple isoforms of cancer stemness regulator NanogP8 in clinical prostate cancer specimens and their unique biochemical and oncogenic properties

**DOI:** 10.64898/2026.11.18.767584

**Authors:** Bigang Liu, Rahul Kumar, Xiaozhuo Liu, Shan Wu, Jianjun Shen, Qiuhui Li, Xin Chen, Kiera Rycaj, Jie Xiong, Ruifang Liu, Rashid Mehmood, Collene Jeter, Dean G. Tang

**Affiliations:** Department of Epigenetics and Molecular Carcinogenesis, the University of Texas M.D Anderson Cancer Center, 1515 Holcombe Blvd., Houston, TX 77030, USA; Department of Pharmacology & Therapeutics, Roswell Park Comprehensive Cancer Center, Elm & Carlton Streets, Buffalo, NY 14263, USA; Experimental Therapeutics (ET) Graduate Program, the University at Buffalo and Roswell Park Comprehensive Cancer Center, Elm & Carlton Streets, Buffalo, NY 14263, USA; The CCSG Developmental Therapeutics (DT) Program, Roswell Park Comprehensive Cancer Center, Elm & Carlton Streets, Buffalo, NY 14263, USA

**Keywords:** Nanog, NanogP8, single nucleotide polymorphism (SNP), prostate cancer, cancer stem cells

## Abstract

The pluripotency regulator Nanog (NANOG, for human protein) has been reported to be overexpressed in cancers and in cancer stem cells (CSCs). We have previously shown that NanogP8, encoded by one of the expressed Nanog pseudogenes, is the predominant transcript expressed in several cancers including prostate cancer (PCa). We have also demonstrated that NanogP8 is functionally important in tumorigenesis as its loss impairs tumor regeneration and growth while inducible expression of NanogP8 is sufficient to reprogram androgen-dependent PCa to an androgen-independent and castration-resistant state. In the current study, we combined RT-PCR, subcloning and sequencing to systematically analyze the open reading frame sequences of *Nanog* and *NanogP8* mRNA transcripts in treatment-naïve clinical human PCa (HPCa) specimens. Our results revealed that HPCa did not express the ES cell-specific *Nanog1* mRNA but primarily expressed *NanogP8* together with several other Nanog pseudogenes including *NanogP7* and *NanogP6*. Interestingly, most HPCa specimens contained multiple isoforms of *NanogP8* transcripts, and we identified novel missense mutations in the *NanogP8* transcripts that could lead to amino acid changes. Experiments with recombinant proteins, site-directed mutagenesis and Western blotting revealed several mutations that conferred the NanogP8 variant proteins, including NanogP8^L61P^, NanogP8^D64Y^, and NanogP8^N130S^, with unique biochemical properties. Moreover, some mutations furnish NanogP8 isoforms with distinct CSC and oncogenic characteristics. Altogether, this study advances our knowledge of the molecular, biochemical and oncogenic properties of Nanog expression and NanogP8 isoforms in PCa.

**Graphical Abstract:** 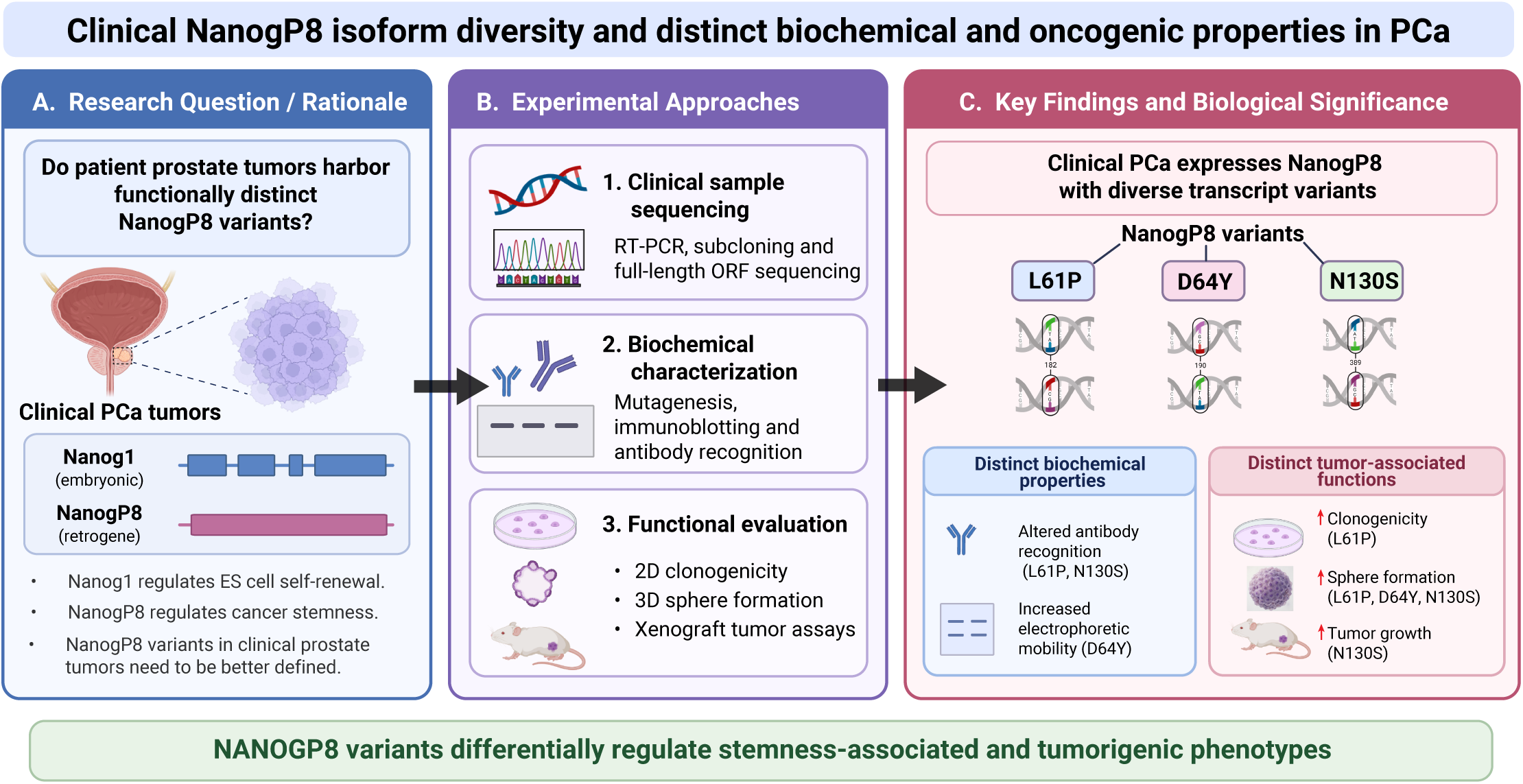

## 1. Introduction

Among the many forms of genetic variations in the human genome, single nucleotide polymorphisms (SNPs) represent the most common type [1, 2]. While SNPs are frequently located in non-coding regions, more than 60,000 have been identified within exonic regions, with approximately 93% of all genes harboring at least one SNP [3]. Non-synonymous coding SNPs, which alter the amino acid (aa) sequence of a protein, can have profound phenotypic consequences by affecting protein structure, solubility, stability, and ultimately, functions [4–6]. These alterations diversify protein functionality and have been frequently associated with cancer initiation and progression [7–9]. Prostate cancer (PCa), like many other malignancies, is associated with and results from the gradual accumulation of genetic alterations [10, 11]. Genome-wide association studies (GWAS) have identified numerous SNPs linked to PCa susceptibility [12]. These genetic variants, present in distinct subclonal populations, can act independently or synergistically to confer a selective growth advantage during tumor evolution. Therefore, dissecting the role of individual non-synonymous coding SNPs, particularly in genes implicated in self-renewal and cancer stemness, will advance our understanding of PCa pathogenesis.

The human Nanog (or NANOG) protein, a key regulator of embryonic stem (ES) cell self-renewal and pluripotency, is encoded by the *Nanog* (i.e., *Nanog1*) gene located on chromosome 12p13.31 [13–15] consisting of 4 exons (**Figure 1A**) with a CDS (coding DNA sequence) of 915 bp (or 918 bp including stop codon) (**Figure 1B**). Expression of Nanog is typically downregulated during ES cell differentiation, in concert with other pluripotency-associated factors [13, 15, 16]. During evolution, the *Nanog* gene underwent tandem duplication to form the paralogous Nanog2 (also known as NanogP1; 915 bp), and multiple retrotransposition events gave rise to a large family of Nanog pseudogenes (NanogP2– NanogP11) scattered throughout the genome [17–19] (**Figure 1B**). With the exception of NanogP1 (i.e., Nanog2), most of these pseudogenes lack introns and have accumulated mutations that render them incapable of producing full-length proteins [17–19]. However, NanogP8, located on chromosome 15q24.2, retains an intact open reading frame (ORF) without apparent structural defects, and it possesses protein-coding potential [18] (**Figure 1B**). Many studies have reported Nanog and/or NanogP8 to be overexpressed and to play oncogenic functions in a variety of human cancers [20–40] including PCa from our own studies [41–43].

**Figure 1.**
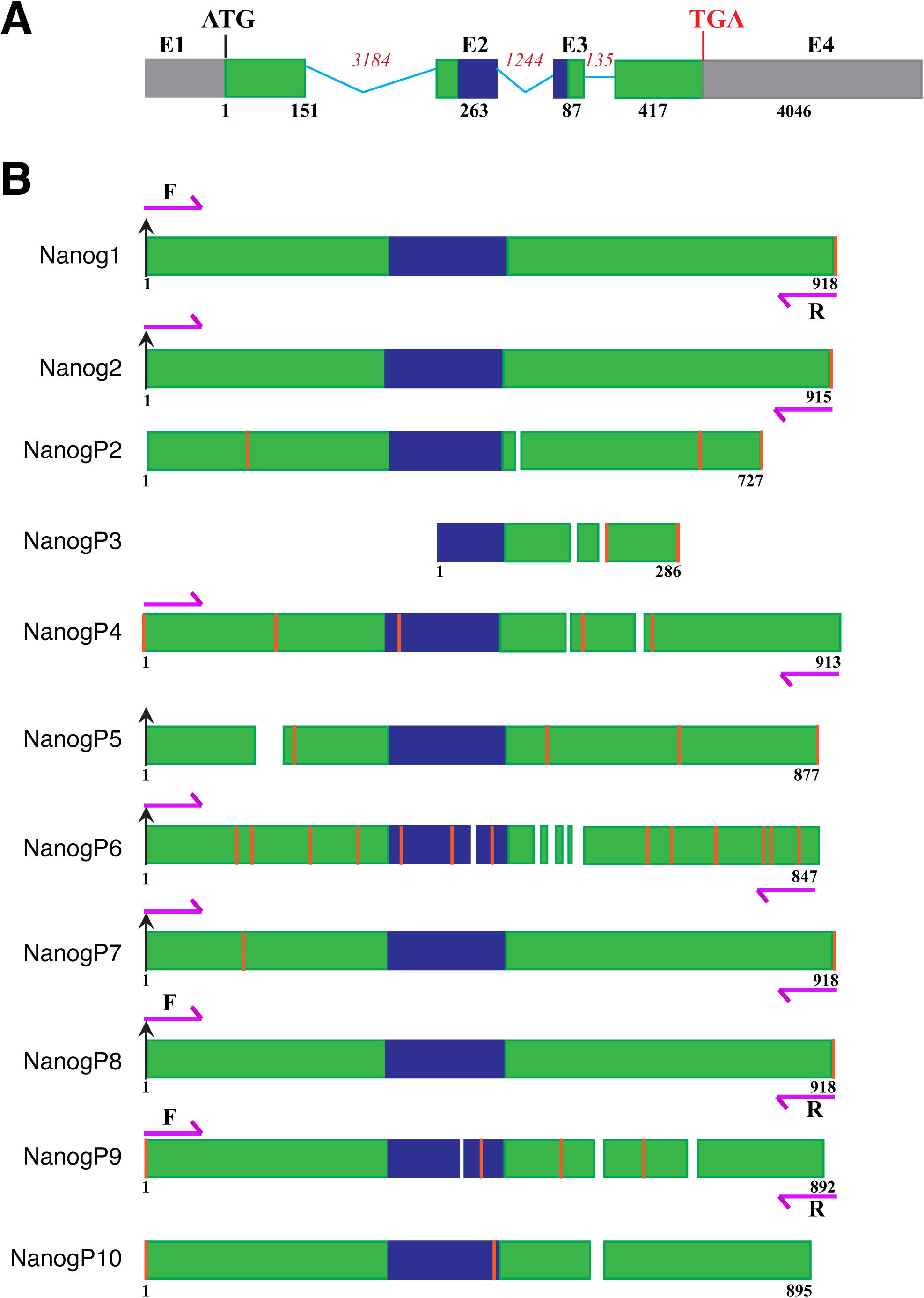
Structure of the Human *Nanog1* gene and Nanog pseudogenes. **(A)** Schematic representation of the human *Nanog1* gene located on chromosome 12p13.31 (based on ENST00000229307.8), which comprises 4 exons (E1 to E4) and 3 introns (marked by blue lines with intron sizes indicated). The green and blue boxes indicate protein-coding regions with blue highlighting the homeobox domain. The grey boxes represent the 5’-and 3’-untranslated regions (UTRs). **(B)** Schematic depiction of the coding regions of *Nanog1* (top) and its processed pseudogenes, along with the primers used for RT-PCR in this study (F, forward; R, reverse). Start codons are indicated by black arrows and stop codons by red vertical lines. Deletions are shown as gaps. Numbers indicate the lengths of the transcripts (in base pairs). The F/R PCR primers can amplify the full-length ORFs of Nanog1, Nanog2 (NanogP1), NanogP4 and NanogP6-NanogP9.

Mechanistically, Nanog/NanogP8 may play a role in maintaining the cancer stem cell (CSC) properties of self-renewal, tumor initiation and propagation, and therapy resistance. Nanog expression enhances these features by activating transcriptional programs linked to stemness, epithelial-to-mesenchymal transition (EMT), and immune evasion [26, 42, 44]. Recent studies have identified subpopulations of metastasis-initiating cells within the prostate and breast tumors that exhibit stem cell-like features and are characterized by elevated expression of CX3CR1 and Nanog [45]. Consistently, work from our laboratory [41–43] and others’ [46–59] has demonstrated a causal relationship between Nanog (or NanogP8) expression and PCa stemness and the progression to castration-resistant PCa (CRPC), a lethal form of the disease that emerges during androgen deprivation therapy (ADT). For example, an immunohistochemical analysis of PCa biopsies from 114 PCa patients revealed that the expression of stem cell markers, including Nanog, was significantly elevated in adenocarcinoma compared to matched benign prostate tissue [48]. Moreover, Nanog expression positively correlated with increasing Gleason scores [48]. Another study involving 85 PCa patients suggested that NanogP8 represents a prognostic factor of PCa progression [49]. Furthermore, treatment of PCa cells with androgen receptor (AR) pathway inhibitors (ARPIs) such as enzalutamide can induce the expression of stemness-associated genes, including Nanog [50, 51]. Nanog has also been implicated in development of resistance to chemotherapeutic agents such as paclitaxel, highlighting its role in treatment failure and disease recurrence. Mechanistically, Nanog may function in cancer cells by interacting with key oncogenic and pro-metastasis signaling regulators and pathways, including Wnt/β-catenin, SPOP, STAT3, and TGF-β [51, 56]. Experimental knockdown of Nanog or inhibition of its downstream signaling impairs tumor growth and sensitizes cancer cells to chemotherapy and hormone therapy [41], suggesting that Nanog is not only a biomarker of aggressive disease but also a viable therapeutic target.

Additionally, Nanog has been implicated in driving immune evasion and immune suppression in several cancers. For instance, Nanog enhances the expression of immunosuppressive molecules such as PD-L1 [60] and facilitates the secretion of anti-inflammatory cytokines including IL-6, IL-10 and CCL5, thereby promoting an immunosuppressive tumor microenvironment [61–63]. This Nanog-modulated tumor immune microenvironment impairs the function of CD8^+^ cytotoxic T cells while promoting the recruitment of regulatory T cells (Tregs), ultimately allowing tumor cells to escape immune surveillance. Furthermore, Nanog may help mediate the resistance to immune checkpoint blockade [60, 64, 65] and promote PCa cells in escaping NK cell attack via downregulating ICAM1 [66]. These studies suggest that therapeutically targeting Nanog or its downstream signaling pathways may enhance immune activation and improve the anti-cancer effectiveness of immunotherapies.

Our group has been systematically studying Nanog expression, regulation, and functions in cancers, with a particular focus on PCa [41–43, 67–72]. For example, we have demonstrated that *NanogP8*, not *Nanog1*, is the predominant transcript expressed in cancer cells [41], and that forced Nanog expression can reprogram PCa cells towards a CSC-like phenotype and a castration-resistant state [42, 43]. Transgenic expression of NanogP8 in the mouse prostate, though insufficient to drive prostate tumorigenesis, promotes development of Hi-Myc tumors [72]. We have also reported several novel mutations and SNPs in *NanogP8* transcripts in somatic cancer cells, some of which seem to endow the NanogP8 protein with unique biochemical properties [68]. Given the importance of NanogP8 in prostate tumorigenesis, a critical question arises: do clinical human PCa (HPCa) samples also harbor SNPs in NanogP8, and if so, what are the functional consequences of the resulting protein isoforms? In the present study, we systematically investigated Nanog expression and mutational profiles in clinical HPCa specimens and PCa cell lines. We confirmed that *NanogP8* mRNA is the predominant Nanog transcript in these samples. Unexpectedly, we also detected transcripts from additional Nanog pseudogenes in clinical specimens. Importantly, we identified multiple novel mutations in *NanogP8* transcripts. Functional analyses revealed that the resulting NanogP8 protein isoforms (or variants) displayed distinct biochemical behaviors, including differential reactivity to commercial antibodies and altered migration patterns on SDS-PAGE gels. Moreover, these NanogP8 isoforms manifested distinct biological properties in PCa cells.

## 2. Results

### 2.1. The HPCa specimens express *NanogP8*, *NanogP7* and *NanogP6* but not *Nanog1* mRNAs

To systematically evaluate the expression of Nanog and its expressed pseudogenes (or retrogenes) and to investigate potential mutation profiles of *NanogP8* mRNAs (**Figure 1**) in patient PCa (i.e., HPCa) specimens, we decided to perform direct analysis of full-length open reading frame (ORF) sequence of *NanogP8* mRNAs from HPCa specimens and PCa cell lines by combining RT-PCR, subcloning and sequencing of the PCR products. We designed a pair of primers that could amplify the ORFs of 7 of the “11 Nanog daughters” ([17–19, 41] including *Nanog1*, *NanogP1, NanogP4,* and *NanogP6 to NanogP9* (**Figure 1B**). Due to significant sequence divergence, this pair of primers would not amplify 4 Nanog pseudogenes, i.e., *NanogP2, NanogP3, NanogP5* and *NanogP10* (**Figure 1B**).

To first confirm our early findings that HPCa cells do not express *Nanog1* mRNA, and instead preferentially express *NanogP8* mRNA [41, 42], we took advantage of a specific mutation [18] at nucleotide (nt) 144 (i.e., nt144G→A) in the exon 1 (E1) of *NanogP8* mRNA that introduces an AlwN1 restriction enzyme site (5’-**CAG**NNN**CTG**-3’) (**Figure 2A**). Thereby, AlwN1 can digest the PCR products from *NanogP8* mRNA but not *Nanog1* mRNA (**Figure 2A**, bottom). We designed a pair of PCR primers flanking the AlwN1 restriction site in the *NanogP8* mRNA (**Figure 2A**). Using this primer pair, we PCR-amplified the 383 bp Nanog1/NanogP8 cDNA fragments from PCa cells derived from cultures (LNCaP, Du145, PC3), xenografts (LAPC4, LAPC9), and a primary patient tumor (HPCa157) as well as 3 lung cancer cell lines (A549, H460, and H1299). Consistent with previous findings [41], we observed that all cancer samples primarily expressed the *NanogP8* mRNA and not *Nanog1* as evidenced by restriction DNA fragments of 261 bp and 122 bp denoting the presence of AlwNI site (**Figure 2B** and **2C**). In contrast, no AlwN1 restriction DNA fragments were observed in the PCR products from the embryonal carcinoma NTERA-2 (N-tera) cells, indicating that these cells solely expressed the *Nanog1* mRNA (**Figure 2B** and **2C**).

**Figure 2.**
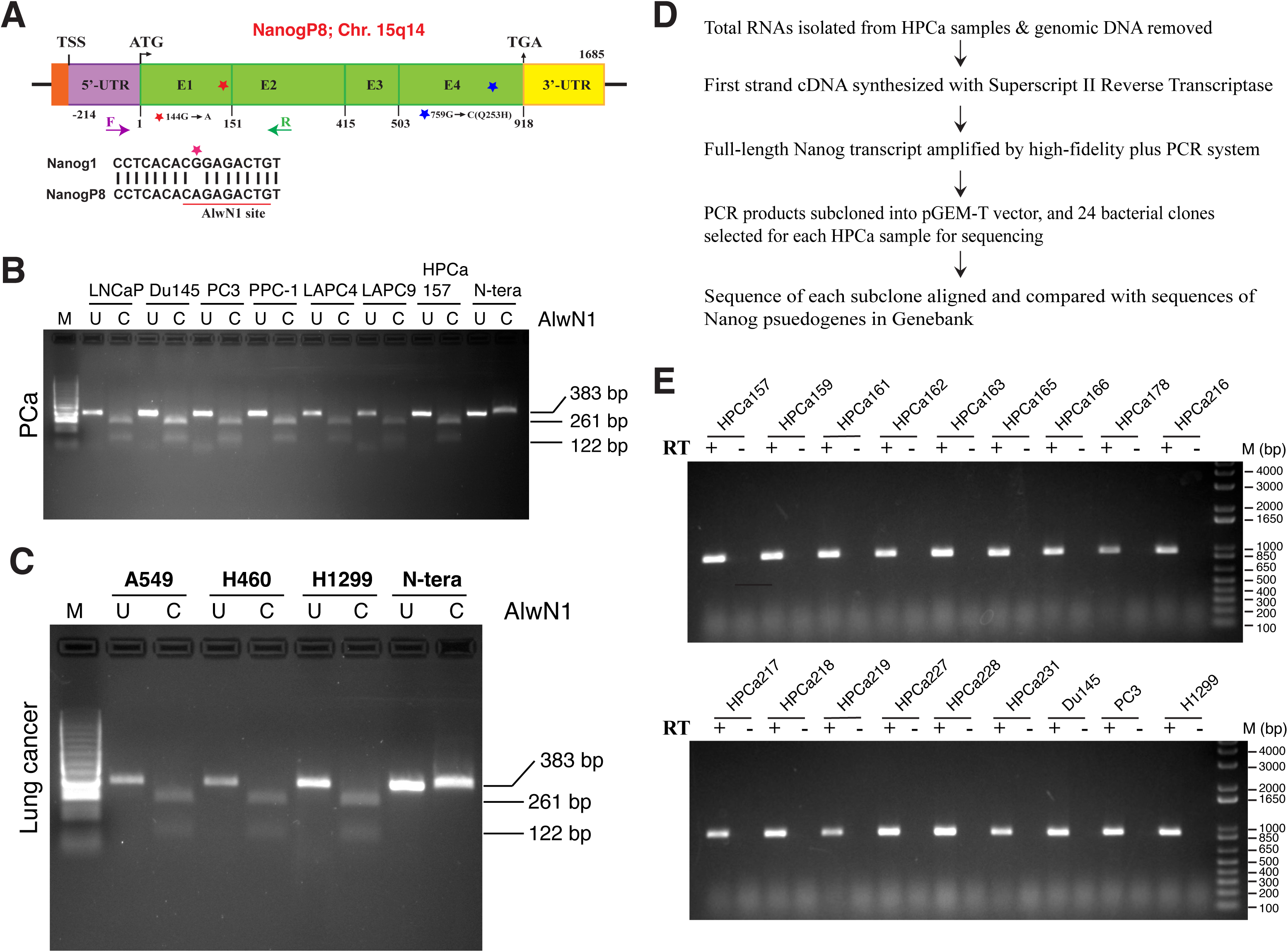
Somatic cancer cells primarily express *NanogP8*. **(A)** Schematic representation of the *NanogP8* mRNA transcript showing the introduction of AlwN1 site (bottom) due to 144G→A mutation in exon (E) 1 (indicated by red *). The TSS (transcriptional start site), ATG (start codon) and TGA (stop codon) as well as the 5’-UTR and 3’-UTR are indicated on top. The pair of PCR primers used to amplify the AlwN1 site-containing region is indicated below. F primer is located in the 5’-UTR (5’-TTGATACTTTTCCTTCTGGAGGTCC-3’, at nucleotide -121 to -96 relative to ATG) and R primer is located in exon 2 (5’-TCCTTTTTTGCGACAC TATTCTCTG-3’; at +238 to +263). Note that the nt759G→C mutation in E4 is predicted to cause Q253H change in aa sequence (indicated by blue *). **(B-C)** mRNAs extracted from the indicated HPCa157 and PCa cell lines (LNCaP, Du145, PC3 and PPC-1) and xenografts (LAPC4, LAPC9) (B), lung cancer cell lines (A549, H460, and H1299) (C), or from the Nanog1 control cell line N-tera were subjected to RT-PCR using the above-mentioned AlwN1-covering PCR primers. The resulting 383-bp PCR products were digested with the restriction enzyme AlwN1, producing two fragments of 261 bp and 122 bp, respectively. Note that only *NanogP8* transcripts due to the characteristic G144A nucleotide substitution are cleavable by AlwN1 whereas *Nanog1* transcripts from N-tera remain undigested. **(D)** Workflow to analyze the expression of full-length *Nanog* isoform transcripts in HPCa and other cancer cells. **(E)** Full-length *Nanog* transcript was amplified by RT-PCR using total RNA isolated from the indicated samples as template. The absence of PCR amplicon in no-RT groups confirmed the complete digestion of genomic DNA.

We then amplified the full-length *Nanog* transcripts, using the primer pair depicted in **Figure 1B**, from 15 freshly collected HPCa samples and 3 cancer cell lines including Du145, PC3 and H1299 (i.e., a total of 18 samples in this study) **Figure 2D** and **2E**). After PCR amplification, we subcloned the PCR products into pGEM-T vector and sequenced 24 clones for each sample from both 5’ and 3’ ends using T7 and SP6 primers (**Figure 2D**). The absence of PCR bands in no-RT controls indicated no contamination of genomic DNA in RT-PCR reactions (**Figure 2E**; lanes labeled “-“). The sequencing results showed that all 15 HPCa samples and the 3 cancer cell lines expressed *NanogP8* mRNA (from 12.5% to 95.8% of the total sequenced transcripts/sample), along with *NanogP4*, *NanogP6*, *NanogP7* and *NanogP9* transcripts at varying levels (**Figure 3A** and **3B**; **Table 1**). We did not detect any *Nanog1* and *Nanog2* (*NanogP1*) mRNA transcripts in any of the samples analyzed (**Table 1**). Interestingly, *NanogP6* mRNA was detected, at 4.3% to 50%, in 11 out of the 15 HPCa samples (i.e., 73%) and the lung cancer cell line H1299 whereas *NanogP4* and *NanogP9* were both detected in 2 of the 18 total samples analyzed (**Figure 3A** and **3B**; **Table 1**). Strikingly, *NanogP7* mRNA was expressed in all examined HPCa samples and cancer cell lines (from 4.2% to 83.3%; **Table 1**). Nevertheless, consistent with the prior knowledge (**Figure 1B**), the *NanogP7* mRNA species detected in all samples contained a stop codon at nt187 (**Figure S1A** and **S1B**).

**Figure 3.**
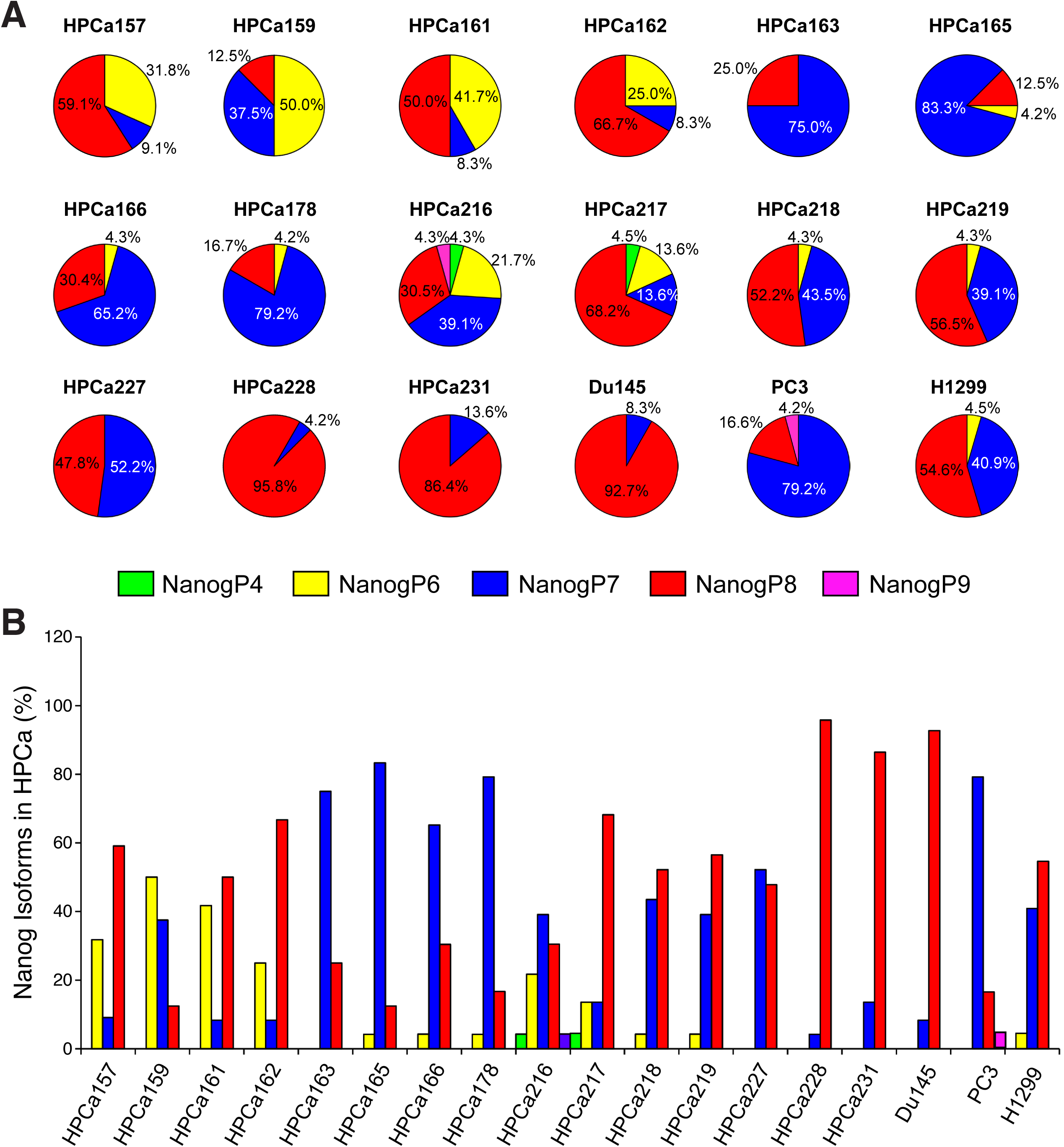
Expression of *Nanog* isoforms in human PCa specimens and cells. **(A-B)** Pie charts (A) and bar graphs (B) presenting percentage of different *Nanog* isoforms including *NanogP4* and *NanogP6-P9* in HPCa samples and indicated cancer cells.

**Table 1.** Expression of Nanog isoforms in human PCa (HPCa) patient samples.

| **Sample No.** | **Gleason Score** | **Nanog1 %** | **Nanog2 %** | **NanogP4 %** | **NanogP6 %** | **NanogP7 %** | **NanogP8 %** | **NanogP9 %** |
|---|---|---|---|---|---|---|---|---|
| HPCa157 | 7 (4+3) | 0 | 0 | 0 | 31.8 | 9.1 | 59.1 | 0 |
| HPCa159 | 6 | 0 | 0 | 0 | 50 | 37.5 | 12.5 | 0 |
| HPCa161 | 9 (4+5) | 0 | 0 | 0 | 41.7 | 8.3 | 50 | 0 |
| HPCa162 | 7 (4+3) | 0 | 0 | 0 | 25 | 8.3 | 66.7 | 0 |
| HPCa163 | 7 | 0 | 0 | 0 | 0 | 75 | 25 | 0 |
| HPCa165 | 7 | 0 | 0 | 0 | 4.2 | 83.3 | 12.5 | 0 |
| HPCa166 | 7 (4+3) | 0 | 0 | 0 | 4.3 | 65.2 | 30.4 | 0 |
| HPCa178 | 9 (5+4) | 0 | 0 | 0 | 4.2 | 79.2 | 16.7 | 0 |
| HPCa216 | 7 (3+4) | 0 | 0 | 4.3 | 21.7 | 39.1 | 30.5 | 4.3 |
| HPCa217 | 7 (3+4) | 0 | 0 | 4.5 | 13.6 | 13.6 | 68.2 | 0 |
| HPCa218 | 7 (4+3) | 0 | 0 | 0 | 4.3 | 43.5 | 52.2 | 0 |
| HPCa219 | 7 (4+3) | 0 | 0 | 0 | 4.3 | 39.1 | 56.5 | 0 |
| HPCa227 | 7 (3+4) | 0 | 0 | 0 | 0 | 52.2 | 47.8 | 0 |
| HPCa228 | 7 (4+3) | 0 | 0 | 0 | 0 | 4.2 | 95.8 | 0 |
| HPCa231 | 7 (4+3) | 0 | 0 | 0 | 0 | 13.6 | 86.4 | 0 |
| Du145 |  | 0 | 0 | 0 | 0 | 8.3 | 92.7 | 0 |
| PC3 |  | 0 | 0 | 0 | 0 | 79.2 | 16.6 | 4.2 |
| H1299 | n/a | 0 | 0 | 0 | 4.5 | 40.9 | 54.6 | 0 |

This Table summarizes the Nanog transcripts detected in our sequencing analysis of 15 HPCa samples and 3 cancer cell lines (bottom). In total, 24 subclones from each sample underwent sequencing analysis (see Methods). The % of a specific Nanog isoform transcript (out of a total of 24 subclones sequenced) is indicated.

### 2.2. Identification of novel single-nucleotide polymorphisms (SNPs) and missense mutations in *NanogP8* transcripts in HPCa specimens

Among all Nanog pseudogenes we detected in HPCa samples, only *NanogP8* has the potential to produce a functional protein. Therefore, we further focused our sequencing analysis on *NanogP8* transcripts. The *NanogP8* mRNA shares over 99% identity to *Nanog1* and has been reported to have 2 fixed diagnostic nucleotide (nt) substitutions in its ORF, i.e., the synonymous (silent) **nt144G→A** change that introduces the AlwN1 site and non-synonymous **nt759G→C** mutation that introduces the Q253H change alongside structural differences in its untranslated regions (UTRs) [18] (**Figure 2A**; indicated by asterisks). In both our previous sequencing of 3 HPCa samples (i.e., HPCa1, 5, 6) and 2 cancer (MCF7 breast cancer and LNCaP PCa) cell lines [41] (**Figure 4A**) as well as the present sequencing analysis in 15 HPCa and 3 cancer cell lines (Du145, PC3 and H1299) (**Table 2**), we detected, in all sample, the nt144G→A and nt759G→C, the two diagnostic signature nt changes. In addition, several other ‘conserved’ nt changes in *NanogP8* have also been reported [17–19] including nt47C→A (A16E mutation), nt164C→T (silent mutation), nt246G→T (K82N mutation), and nt531T→C (silent mutation), all of which were detected in our studies (**Figure 4A**; **Table 2**). Taken together our early [41] and current studies, we considered the *NanogP8* mRNA with the 4 conserved mutations, i.e., nt144G→A, nt246G→T, nt531T→C, and nt759G→C (**Table 2**), as the ‘wild-type’ NanogP8 (**NanogP8^WT^**).

**Figure 4.**
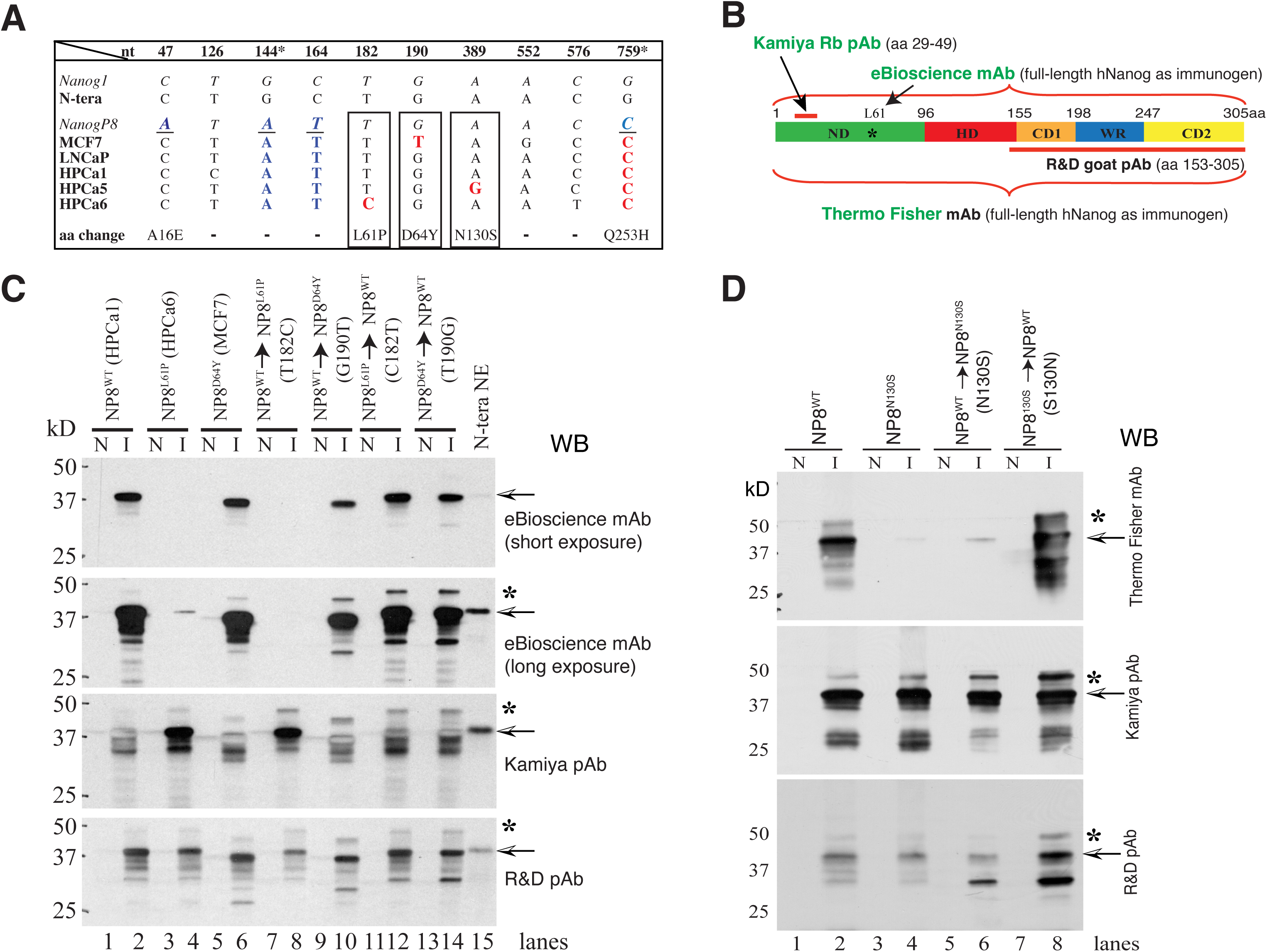
Site-directed mutagenesis reveals distinct biochemical properties of NanogP8 proteins. **(A)** Predicted amino acid (aa) differences between Nanog1 and NanogP8 based on our early cDNA sequencing results [41]. The reported *NanogP8* cDNA (the third row) contains at least 4 conserved nucleotide (nt) differences (marked in bold and underlined blue italics) compared to *Nanog1* cDNA (top row), including the two fixed nt changes (indicated by * at the top). Some of the conserved nt changes in *NanogP8* result in aa substitutions (e.g., Q253H) indicated at the bottom. Three additional nt differences unique to the *NanogP8* transcripts in HPCa6 (T182C), MCF7 (G190T), and HPCa5 (A389G), respectively, were boxed and further studied at protein levels. **(B)** Schematic representation of epitopes recognized by various anti-Nanog antibodies (Abs; pAb, polyclonal Ab; mAb, monoclonal Ab) used in this study. **(C–D)** Western blot (WB) analysis of Nanog1 protein and NanogP8 protein variants. Arrows mark the 42 kD NanogP8 protein [68] and stars indicate minor higher molecular weight bands. Antibodies used are presented on the right and molecular weight markers are shown on the left. Both short and long exposures are provided for the eBioscience mAb (C). Nanog1 protein expressed in N-tera cells serves as the positive control. Abbreviations: NP8, NanogP8; I, induced by IPTG; N, non-induced.

**Table 2.**
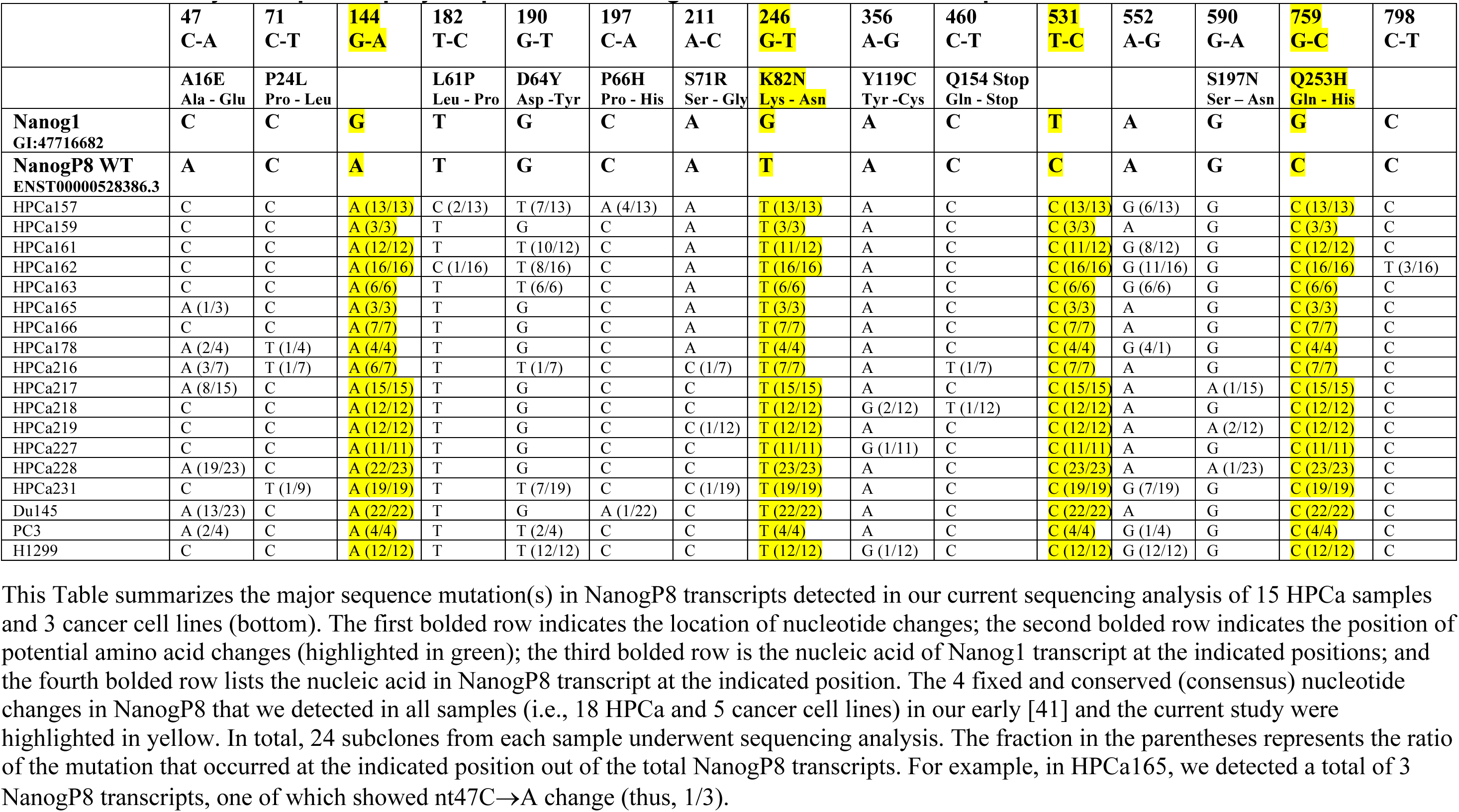
Summary of sequence polymorphisms in NanogP8 isoforms in HPCa samples*.

|  | **47<br>C-A** | **71<br>C-T** | **144<br>G-A** | **182<br>T-C** | **190<br>G-T** | **197<br>C-A** | **211<br>A-C** | **246<br>G-T** | **356<br>A-G** | **460<br>C-T** | **531<br>T-C** | **552<br>A-G** | **590<br>G-A** | **759<br>G-C** | **798<br>C-T** |
|  | **A16E<br>Ala - Glu** | **P24L<br>Pro - Leu** |  | **L61P<br>Leu - Pro** | **D64Y<br>Asp -Tyr** | **P66H<br>Pro - His** | **S71R<br>Ser - Gly** | **K82N<br>Lys - Asn** | **Y119C<br>Tyr -Cys** | **Q154 Stop<br>Gln - Stop** |  |  | **S197N<br>Ser – Asn** | **Q253H<br>Gln - His** |  |
|---|---|---|---|---|---|---|---|---|---|---|---|---|---|---|---|
| **Nanog1<br>GI:47716682** | **C** | **C** | **G** | **T** | **G** | **C** | **A** | **G** | **A** | **C** | **T** | **A** | **G** | **G** | **C** |
| **NanogP8 WT<br>ENST00000528386.3** | **A** | **C** | **A** | **T** | **G** | **C** | **A** | **T** | **A** | **C** | **C** | **A** | **G** | **C** | **C** |
| HPCa157 | C | C | A (13/13) | C (2/13) | T (7/13) | A (4/13) | A | T (13/13) | A | C | C (13/13) | G (6/13) | G | C (13/13) | C |
| HPCa159 | C | C | A (3/3) | T | G | C | A | T (3/3) | A | C | C (3/3) | A | G | C (3/3) | C |
| HPCa161 | C | C | A (12/12) | T | T (10/12) | C | A | T (11/12) | A | C | C (11/12) | G (8/12) | G | C (12/12) | C |
| HPCa162 | C | C | A (16/16) | C (1/16) | T (8/16) | C | A | T (16/16) | A | C | C (16/16) | G (11/16) | G | C (16/16) | T (3/16) |
| HPCa163 | C | C | A (6/6) | T | T (6/6) | C | A | T (6/6) | A | C | C (6/6) | G (6/6) | G | C (6/6) | C |
| HPCa165 | A (1/3) | C | A (3/3) | T | G | C | A | T (3/3) | A | C | C (3/3) | A | G | C (3/3) | C |
| HPCa166 | C | C | A (7/7) | T | G | C | A | T (7/7) | A | C | C (7/7) | A | G | C (7/7) | C |
| HPCa178 | A (2/4) | T (1/4) | A (4/4) | T | G | C | A | T (4/4) | A | C | C (4/4) | G (4/1) | G | C (4/4) | C |
| HPCa216 | A (3/7) | T (1/7) | A (6/7) | T | T (1/7) | C | C (1/7) | T (7/7) | A | T (1/7) | C (7/7) | A | G | C (7/7) | C |
| HPCa217 | A (8/15) | C | A (15/15) | T | G | C | C | T (15/15) | A | C | C (15/15) | A | A (1/15) | C (15/15) | C |
| HPCa218 | C | C | A (12/12) | T | G | C | C | T (12/12) | G (2/12) | T (1/12) | C (12/12) | A | G | C (12/12) | C |
| HPCa219 | C | C | A (12/12) | T | G | C | C (1/12) | T (12/12) | A | C | C (12/12) | A | A (2/12) | C (12/12) | C |
| HPCa227 | C | C | A (11/11) | T | G | C | C | T (11/11) | G (1/11) | C | C (11/11) | A | G | C (11/11) | C |
| HPCa228 | A (19/23) | C | A (22/23) | T | G | C | C | T (23/23) | A | C | C (23/23) | A | A (1/23) | C (23/23) | C |
| HPCa231 | C | T (1/9) | A (19/19) | T | T (7/19) | C | C (1/19) | T (19/19) | A | C | C (19/19) | G (7/19) | G | C (19/19) | C |
| Du145 | A (13/23) | C | A (22/22) | T | G | A (1/22) | C | T (22/22) | A | C | C (22/22) | A | G | C (22/22) | C |
| PC3 | A (2/4) | C | A (4/4) | T | T (2/4) | C | C | T (4/4) | A | C | C (4/4) | G (1/4) | G | C (4/4) | C |
| H1299 | C | C | A (12/12) | T | T (12/12) | C | C | T (12/12) | G (1/12) | C | C (12/12) | G (12/12) | G | C (12/12) | C |

This Table summarizes the major sequence mutation(s) in NanogP8 transcripts detected in our current sequencing analysis of 15 HPCa samples and 3 cancer cell lines (bottom). The first bolded row indicates the location of nucleotide changes; the second bolded row indicates the position of potential amino acid changes (highlighted in green); the third bolded row is the nucleic acid of Nanog1 transcript at the indicated positions; and the fourth bolded row lists the nucleic acid in NanogP8 transcript at the indicated position. The 4 fixed and conserved (consensus) nucleotide changes in NanogP8 that we detected in all samples (i.e., 18 HPCa and 5 cancer cell lines) in our early [41] and the current study were highlighted in yellow. In total, 24 subclones from each sample underwent sequencing analysis. The fraction in the parentheses represents the ratio of the mutation that occurred at the indicated position out of the total NanogP8 transcripts. For example, in HPCa165, we detected a total of 3 NanogP8 transcripts, one of which showed nt47C→A change (thus, 1/3).

Several perspectives were obtained from our current large-scale sequencing efforts. *FIRST*, in addition to above-mentioned 4 highly conserved nt changes, we detected a total of 148 mutations/SNPs in *NanogP8* mRNAs, ∼65% of which could result in aa changes with 9 mutations introducing premature stop codons (**Table 2** and **Table S1**). *SECOND*, most HPCa samples (80%) contained *NanogP8* transcripts with all 4 conserved mutations but only HPCa159 expressed all *NanogP8* transcripts as NanogP8^WT^ without additional variants while the *NanogP8* transcripts from the majority of HPCa specimens showed multiple mutations (1 to 13) (**Table S1**). *THIRD*, consistent with our previous report [41], the reported ‘conserved’ nt47C→A (A16E) mutation [18] was detected in only some transcripts from 5 HPCa specimens (from 33.33% to 82.61%), and Du145 (59.09%) and PC3 (50.00%) cells (**Table 2**). Most HPCa did not display this mutation, suggesting that the nt47C→A mutation is not well conserved in PCa. *FINALLY*, of great interest, some novel *NanogP8* mRNA variants appeared to be recurrent in HPCa. For example, multiple *NanogP8* transcripts from HPCa157, 161, 162, 163, 216 and 231 contained the nt190G→T mutation, which we reported before in MCF7 cells [41] and would introduce the D64Y aa change in NanogP8 protein, i.e., **NanogP8^D64Y^** (**Figure 4A**; **Table 2**). Strikingly, all *NanogP8* transcripts from HPCa163 and the lung cancer cell line H1299 had this mutation (**Table 2** and **Table S1**). Also, several *NanogP8* transcripts from both HPCa157 and HPCa162 not only contained the nt190G→T mutation (i.e., the NanogP8^D64Y^ variant) but also nt182T→C mutation (**Table 2** and **Table S1**), which would induce the L61P aa change, i.e., **NanogP8^L61P^**, which we previously identified in HPCa6 (**Figure 4A**). Finally, we observed the nt389A>G mutation in HPCa5 [41], which could introduce N130S aa change (i.e., **NanogP8^N130S^**) (**Figure 4A**). As these three mutations, D64Y, L61P, and N130S, could represent “hot-spot” mutations in *NanogP8* mRNA in PCa, we investigated their biochemical and biological properties in the following studies.

### 2.3. NanogP8 protein isoforms exhibit unique and isoform-intrinsic biochemical properties

As the NanogP8^L61P^ and NanogP8^D64Y^ mutations are abundant in patient HPCa (above), we first studied their biochemical behavior, together with NanogP8^N130S^ (**Figure 4A**; **Figure S2** and **Figure S3**). We subcloned the full-length cDNAs of NanogP8^WT^ (from HPCa1) NanogP8^L61P^ (from HPCa6), NanogP8^D64Y^ (from MCF7) and NanogP8^N130S^ (from HPCa5) (**Figure 4A**) into lentiviral vector pLVX-IRES-Zsgreen1 to generate NanogP8 lentiviral expression constructs (**Figure S2A-B**). For comparison, we used *Nanog1* cDNA derived from NTERA-2 cells to express Nanog1 protein [68]. We first examined the localization of NanogP8 protein species by transducing LNCaP and Du145 cells with various NanogP8-encoding lentiviral vectors (**Figure S3A-B**). Immunostaining with an anti-Nanog antibody (Ab) revealed that all NanogP8 species, like Nanog1, displayed a similar nuclear localization pattern (**Figure S3A-B**), a typical subcellular localization expected for a transcription factor. These results suggest that individual mutations/SNPs in NanogP8 protein did not affect the nuclear localization of Nanog proteins in PCa cells.

We then tested, by Western blotting (WB), the reactivities of Nanog1 and NanogP8 species to 5 commercial anti-Nanog Abs raised against the human Nanog1 protein (**Figure 4B** and **Figure S2A**). Among the five anti-Nanog Abs, two (Kamiya Rb pAb and Cell Signaling (CS) Rb mAb) were directed against the N-terminus of Nanog protein, one (R&D goat pAb) to the C-terminus, whereas the other two monoclonal Abs (eBiosciences and Thermo Fisher mAbs) were raised against the full-length human Nanog1 protein (**Figure 4B**). We infected the HEK293 cells with lentiviral vectors encoding Nanog1 or NanogP8 isoforms and, as shown in **Figure S2A-B**, different anti-Nanog Abs exhibited distinct reactivities to the five Nanog/NanogP8 protein species. *First*, all Abs detected an obvious ∼42 kD band, previously identified as the dominant Nanog protein band in both NTERA-2 and somatic cancer cells [68], and a band of ∼35 kD (the predicted Nanog protein molecular weight) as well as several additional bands ranging from 25 kD to 40 kD (**Figure S2A-B**). *Second*, different Nanog/NanogP8 protein species displayed differential reactivity to anti-Nanog Abs with slightly different patterns. Three Abs, CS mAb, R&D pAb, and Kamiya mAb, detected all five Nanog/NanogP8 protein species with overall similar efficiency (**Figure S2A**). However, two mAbs, namely, the eBioscience mAb and Thermofisher mAb, did not recognize NanogP8^L61P^ and NanogP8^N130S^, respectively (**Figure S2B**). *Third*, the NanogP8^D64Y^ protein consistently migrated slightly faster, in all WB with the five antibodies, than other Nanog species (**Figure S2A-B**), much like our previous findings where NanogP8 protein species were expressed in bacteria [68]. The above results suggest that NanogP8 proteins with specific mutations/SNPs may confer unique and isoform-intrinsic biochemical properties that allow different NanogP8 protein species to adopt distinct conformation, thus showing unique reactivities (than Nanog1) to different anti-Nanog Abs. To further investigate the two striking observations, i.e., NanogP8^L61P^ and NanogP8^N130S^ proteins could not be recognized by eBioscience and Thermofisher anti-Nanog Abs, respectively and the NanogP8^D64Y^ protein migrated slightly faster than Nanog1 and other NanogP8 proteins, we performed site-directed mutagenesis and expressed these recombinant Nanog (rNanog) proteins in bacteria. As shown in **Figure 4C** and **4D**, the rNanogP8^L61P^ protein was not recognized by the eBiosceince mAb and rNanogP8^N130S^ did not react with Thermofisher mAb whereas rNanogP8^D64Y^ protein migrated faster than NanogP8^WT^ and endogenous Nanog1 protein in NTERA-2 cells. Remarkably, when rNanogP8^WT^ was changed to rNanogP8^L61P^ by mutating nt182T→C (**Figure 4A**), it was no longer recognized by the eBioscience mAb (**Figure 4C**, lane 8). Similarly, when the rNanogP8^WT^ was changed to rNanogP8^D64Y^ by mutating nt190G→T, the protein now showed increased mobility (**Figure 4C**, lane 10). Moreover, rNanogP8^WT^ could not react with the Thermofisher mAb after mutating the nt389A→G (**Figure 4D**, lane 6). Consistently, when we performed the reverse mutagenesis experiments by changing rNanogP8^L61P^, rNanogP8^D64Y^, and rNanogP8^N130S^ proteins to rNanogP8^WT^, these changes not only restored the reactivities of rNanogP8^L61P^ and rNanogP8^N130S^ proteins to eBioscience and Thermofisher mAbs, respectively (**Figure 4C**, lane 12; **Figure 4D**, lane 8), but also reinstated the normal mobility of rNanogP8^D64Y^ (**Figure 4C**, lane 14).

We reported previously that the Kamiya pAb preferentially recognized the rNanogP8^L61P^ whereas the R&D pAb recognized all rNanog proteins equally well [68]. These observations were corroborated herein in the site-directed mutagenesis experiments (**Figure 4C-D**). Collectively, these results reveal two critical points: first, single aa changes can fundamentally affect the biochemical properties and change the conformations of NanogP8 proteins, and second, different anti-Nanog Abs may recognize NanogP8 proteins of different conformations with varying efficiencies.

### 2.4. Nanogp8 isoforms exhibit differential PCa-promoting effects in vitro and in vivo

Our previous studies indicate that 1) endogenous NanogP8 plays a role in promoting the CSC and tumorigenic properties in multiple cancer cell types including PCa [41] and 2) inducible NanogP8 expression is sufficient to reprogram androgen-dependent PCa to a castration-resistant state in cell and xenograft models [42, 43]. Since L61P, D64Y and N130S NanogP8 mutations are identified in patient HPCa specimens and likely clinically relevant and these mutations impart differential biochemical properties to NanogP8 proteins (above), we next explored their potential biological significance in PCa cells. We first conducted in vitro two-dimensional (2D) clonal and three-dimensional (3D) sphere formation assays by overexpressing different NanogP8 isoforms in AR^+^ LNCaP (**Figure 5**) and AR^-^Du145 (**Figure 6A**) cells. After infecting LNCaP and Du145 cells with lentiviruses encoding the 4 NanogP8 isoforms (namely, NanogP8^WT^, NanogP8^L61P^, NanogP8^D64Y^, and NanogP8^N130S^), Nanog1 or empty vector (EV) at a multiplicity of infection (MOI) of 20 for 48 h, we examined the expression levels of Nanog proteins by WB analysis and found that all Nanog1/NanogP8 protein species were overexpressed (relative to EV cells) at comparable levels in transduced cells (data not shown).

**Figure 5.**
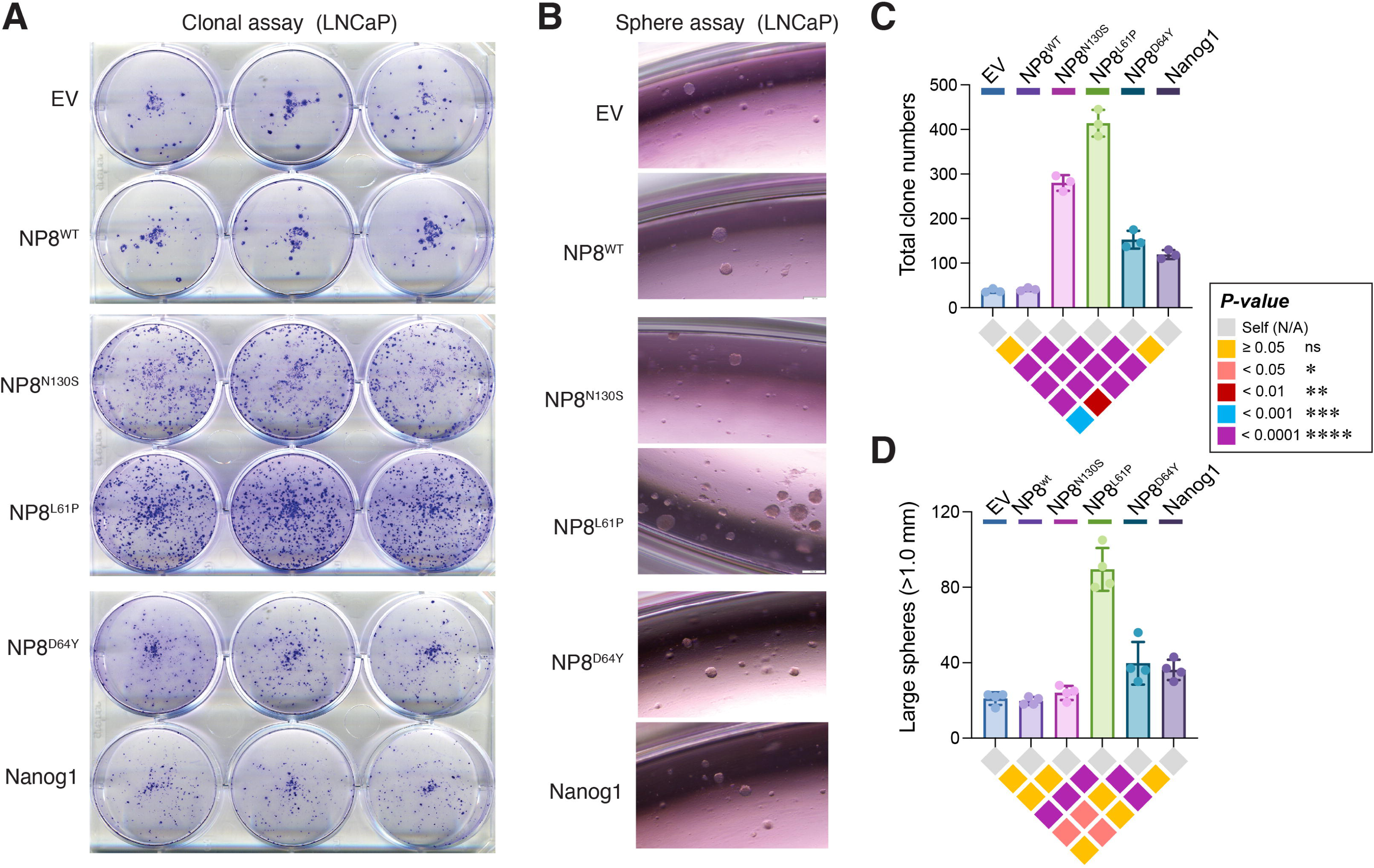
Nanogp8 isoforms display distinct biological activities in LNCaP cells. **(A)** LNCaP cells were infected with indicated lentiviral particles to express the Nanog1, NanogP8 and other NanogP8 mutants for 48 h and cells were trypsinized and seeded in a 6-well plate for 7 days. **(B)** LNCaP cells were infected with the indicated lentiviruses encoding Nanog1, NanogP8 and other NanogP8 mutants for 48 h and cells were trypsinized and mixed with Matrigel and seeded on the edge of a 24-well plate for 14 days. **(C–D)** The total numbers of 2D clones (C; based on A) and large spheres (>1.0 mm in diameter; D, based on B) were quantified and presented as bar graphs (mean ± SD), with individual data points overlaid. Statistical significance was determined using one-way ANOVA followed by Tukey’s multiple-comparisons test (GraphPad Prism 11). All pairwise comparisons among the six groups are visualized using color-coded inverted pyramid significance matrices beneath the bar graphs, with each colored diamond representing a unique pairwise comparison. Colors indicate the ranges of Tukey-adjusted *P*-values, as defined in the accompanying boxed legend (right). Uncolored (gray) diamonds on the top row represent self-comparisons included for visual symmetry.

**Figure 6.**
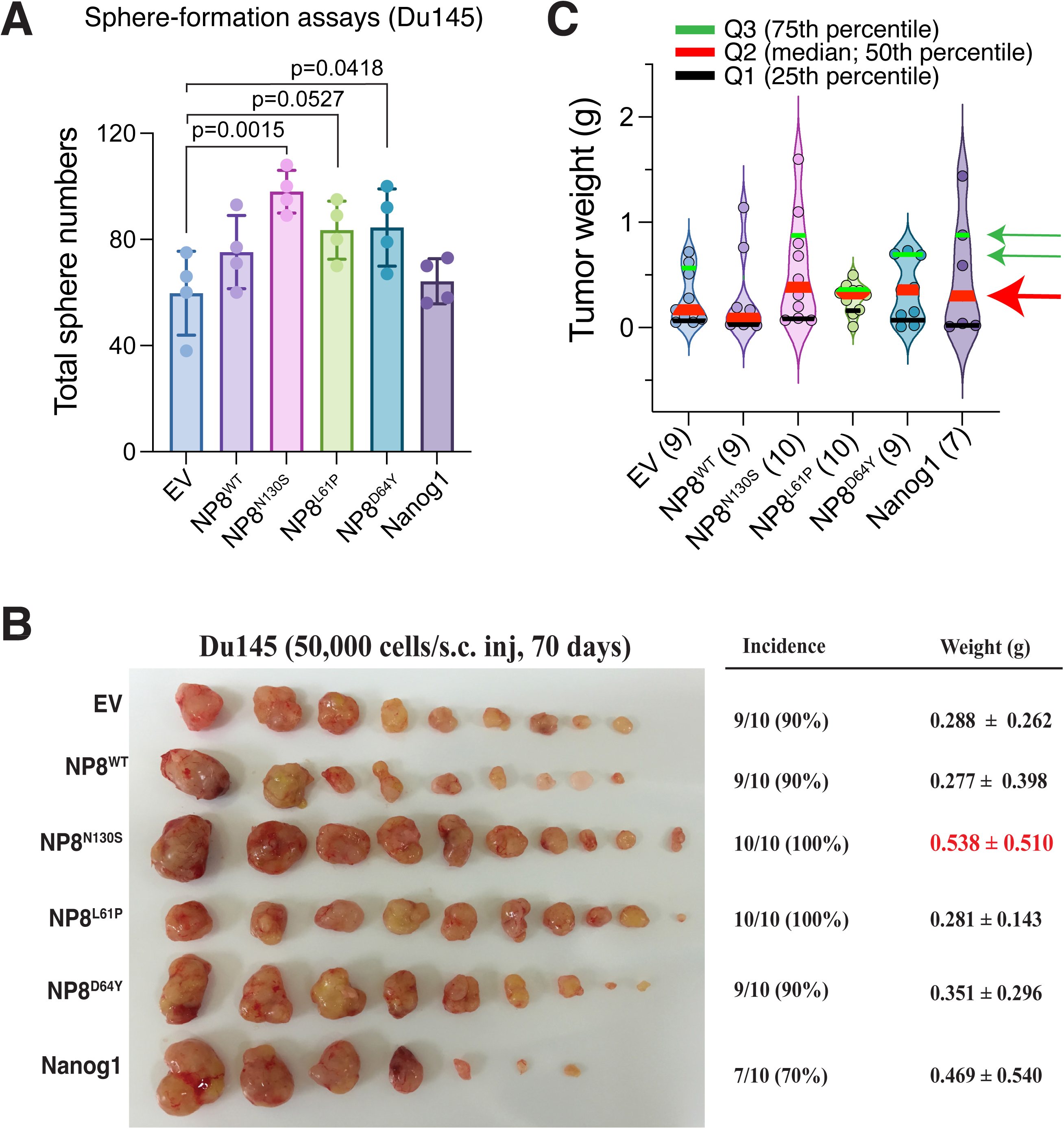
NanogP8 protein isoforms displayed distinct sphere- and tumor-promoting activities in Du145 cells. (**A**) Sphere-formation assays were performed as in LNCaP cells and total number of spheres were presented as bar graphs (mean ± S.D). The 3 NanogP8 variants promoted sphere formation in Du145 cells to varying degrees with NP8^N130S^ mutant showing the strongest effect. Statistical significance was determined using one-way ANOVA followed by Dunnett’s multiple-comparisons test vs. the EV control. (**B**) Overexpression of NP8^N130S^ promoted Du145 tumor growth in vivo. Du145 cells were infected with the indicated lentiviruses encoding the empty vector (EV; control), Nanog1, NanogP8 and other NanogP8 mutants, and subcutaneously (s.c) injected (cell # indicated) in male NOD/SCID mice. After 70 days, mice were sacrificed and tumors harvested. Shown are endpoint tumor images (left), tumor incidence (middle), and tumor weight (mean ± S.D; right). (**C**) The NanogP8 variants, especially the NanogP8^N130S^, promoted generation of large-sized Du145 tumors. Violin plots show the distributions of individual tumor weights from tumor-bearing mice in the indicated groups. Horizontal lines indicate the 25th percentile (Q1, black), median (Q2, red), and 75th percentile (Q3, green). Quartile estimates are summarized descriptively, with exploratory 95% confidence intervals estimated using nonparametric bootstrap resampling (50,000 iterations). Several experimental groups, particularly NanogP8^N130S^, exhibited higher median (Q2) and upper-quartile (Q3) tumor weights than NanogP8^WT^ (or EV) See Text.

In low cell-plating 2D clonal assays [41, 73–75], we observed that overexpression of Nanog1 (but not NanogP8^WT^) increased colony formation (clonality) in LNCaP cells compared to the EV group (**Figure 5A** and **Figure 5C**). Strikingly, overexpression of all three patient HPCa-derived NanogP8 mutants, especially the NanogP8^L61P^ and NanogP8^N130S^, considerably promoted the clonogenicity of LNCaP cells in comparison to NanogP8^WT^ (**Figure 5A** and **Figure 5C**). Next, we tested the effect of these NanogP8 mutants on “stemness” of the manipulated LNCaP and Du145 cells using the 3D sphere-formation assays in Matrigel [11, 68, 69]. The results showed that overexpression of NanogP8^L61P^ in LNCaP cells markedly increased both the total number of spheres (**Figure 5B**) and the number of large (diameter >1.0 mm) spheres (**Figure 5D**). Nanog1 and NanogP8^D64Y^ also increased the abundance of large spheres (**Figure 5B** and **5D**). In Du145 cells, although neither NanogP8^WT^ nor Nanog1 overexpression affected sphere formation compared to the EV group, overexpression of NanogP8^N130S^ and NanogP8^D64Y^ both enhanced sphere formation while NanogP8^L61P^ overexpression showed a statistical trend toward sphere promotion (**Figure 6A**). These in vitro studies, collectively, indicate that the 3 NanogP8 variants, in a cell type-dependent manner and to varying degrees, enhance clonogenic and sphere-forming capabilities in PCa cells.

Finally, we assessed the potential in vivo tumorigenic impacts of Nanog1, NanogP8^WT^ and the 3 NanogP8 variants in Du145 cells. Briefly, 50,000 Du145 cells transduced with various constructs were subcutaneously (s.c) implanted in male NOD/SCID mice and tumors were harvested 70 days post cell implantation. As shown in **Figure 6B**, consistent with the NanogP8^N130S^ promoting Du145 cell sphere formation (**Figure 6A**), overexpression of NanogP8^N130S^ isoform also led to endpoint Du145 tumors approximately twice as large as those in the EV and NanogP8^WT^ groups. Although the averaged (mean) tumor weights in NanogP8^N130S^ group as well as in other NanogP8 variant and Nanog1 groups were not statistically different compared to EV or NanogP8^WT^ group due to wide variations in tumor sizes in each group, it was nevertheless apparent that all 4 experimental groups (i.e., Nanog1 and 3 NanogP8 variants) all had more large-sized tumors (**Figure 6B**). In support, when individual tumors in each group were segregated in violin plots and presented in quartiles (Q), it became clear that although the Q1 (i.e., the lower 25^th^ percentile) tumors in all 6 groups were similar (**Figure 6C**, black lines), the 4 experimental groups, especially the NanogP8^N130S^ group, showed many more Q2 (i.e., the median 50^th^ percentile) tumors (**Figure 6C**, red lines indicated by a red arrow) and more Q3 (i.e., the 75^th^ percentile) tumors (**Figure 6**, green lines marked by green arrows) when compared to the NanogP8^WT^ group. Indeed, the Q2 tumors in NanogP8^WT^, NanogP8^L61P^, NanogP8^D64Y^, NanogP8^N130S^, Nanog1 groups were 0.09g (0.03-0.76g), 0.32g (0.17-0.38g), 0.36g (0.02-0.7g), 0.39g (0.09-0.89g), and 0.3g (0.02-0.88g), respectively (95% confidence intervals). Similarly, the Q3 tumors in NanogP8^WT^, NanogP8^L61P^, NanogP8^D64Y^, NanogP8^N130S^, Nanog1 groups were 0.19g (0.06-1.14g), 0.35g (0.298-0.48g), 0.69g (0.15-0.73g), 0.77g (0.28-1.48g), and 0.74g (0.04-1.44g), respectively (95% confidence intervals).

## 3. Discussion

Our previous studies have shown that **1)** NanogP8 is preferentially expressed in various somatic cancer cells including PCa [41] and (AR^-/lo^)PSA^-/lo^ PCa stem cells [75]; **2)** NanogP8 plays a functional role in conferring cancer-associated properties as its loss or knockdown impairs clonogenic and sphere-forming capabilities as well as tumor regeneration [41]; **3)** inducible expression of NanogP8 is sufficient, in 2-3 weeks, to reprogram androgen-dependent PCa to an androgen-independent and castration-resistant state via dynamically repressing and engaging the AR, FOXA1 and MYC signaling [42, 43]; **4)** high levels of transgenic NanogP8 expression in the mouse K14 cellular compartment disrupt developmental programs in multiple tissues leading to perinatal lethality [67], and **5)** transgenic expression of NanogP8 in the mouse prostate, though insufficient to induce tumor formation, promotes Myc-driven prostate tumorigenesis [72]. Others have also reported preferential expression of NanogP8 in somatic cancer cells [20, 28, 76, 77], NanogP8 knockout impairing tumorigenic and CSC properties [37, 58, 78], and the reprogramming activities of NanogP8 [76].

In the present study, we set out to systematically investigate NanogP8 expression and its polymorphic (mutational) landscape in clinical HPCa specimens and commonly used PCa cell lines. Due to the high sequence homology between Nanog1 and NanogP8, differing by only four conserved nucleotides and potentially two amino acids (i.e., nt144G→A, nt246G→T (K82N), nt531T→C, and nt759G→C (Q253H)), distinguishing between them at the protein level using conventional antibody-based approaches remains challenging. Therefore, we combined clonal sequencing and molecular analysis to characterize the full-length open reading frames of all *Nanog* mRNA species in a total of 18 HPCa samples (including 15 in current study and 3 in the early study ([41] and **Figure 4A**) and 5 cancer (including 3 PCa – LNCaP, Du145 and PC3; 1 breast cancer – MCF7; and 1 lung cancer) cell lines (**Figure 4A**; **Table 2**). We confirmed our previous findings [41] that patient HPCa do not express the ES-specific *Nanog1* but predominantly express the processed retrogene-derived *NanogP8* (**Table 1**). Notably, we identified several novel missense mutations within *NanogP8* transcripts, across tumor samples, many of which introduce amino acid substitutions with potential functional consequences. For example, beyond the canonical differences between NanogP8 and Nanog1 (i.e., K82N, Q253H), we identified several recurrent missense mutations in HPCa samples—including A16E, P24L, L61P, D64Y, and S197N—with most tumors expressing several *NanogP8* isoforms (**Table 2**). Western blotting and site-directed mutagenesis revealed that specific point mutations altered the antigenicity (i.e., antibody reactivity) and biochemical properties of NanogP8 isoforms. For example, the NanogP8^L61P^ isoform was not recognized by an anti-Nanog mAb from eBioscience, while the NanogP8^N130S^ isoform failed to react with an anti-Nanog mAb from Thermofisher. In contrast, both antibodies readily detected NanogP8^WT^ and other isoforms. Moreover, the NanogP8^D64Y^ isoform consistently exhibited slightly faster electrophoretic mobility compared to Nanog1 and other NanogP8 variants. These results extend our earlier studies reporting on the interesting properties of Nanog1 protein in embryonal carcinoma cells and recombinant NanogP8 proteins from somatic cancer cells [68] and suggest that point mutations in NanogP8, identified in patient HPCa samples, may profoundly affect its biochemical properties and modulate the biological functions of NanogP8 variants. Indeed, our prospective gain-of-function studies indicate that different NanogP8 isoforms seem to promote cancer-associated and tumorigenic properties in PCa cells in a cell type-dependent manner. Thus, the NanogP8^L61P^ mutant markedly increased both clonogenic and sphere-founding properties in LNCaP cells compared to NanogP8^WT^ whereas the NanogP8^N130S^ variant enhanced clonality (but not sphere formation) of LNCaP cells and promoted both sphere formation and tumor growth in the Du145 model (**Figure 5** and **Figure 6**).

Based on above findings as well as inherent PCa heterogeneity, it is tempting to speculate that distinct tumor cell subclones (or subtypes) may express different NanogP8 isoforms, which may possess unique biochemical and biological properties that enable PCa cells to adapt to changing environment (e.g., hypoxia) and genetic alterations (e.g., *RB1* loss or *MYC* amplification). Consequently, different tumor cell subpopulations may cooperate to evade the host immune system and collectively drive PCa progression. Moreover, as previously reported by our group and others, Nanog and other pluripotency-associated factors are upregulated in PCa especially during progression and following ADT, potentially promoting the reprogramming of lineage-committed tumor cells into stem cell-like state and contributing to CRPC [41–43, 46–48, 50–52]. Expression of NanogP8 (and Nanog in general) may be regulated at the transcriptional, post-transcriptional, and protein stability levels. For example, NanogP8 expression in PCa has been associated with activation of the 5’-flanking region of the retrogene [24] and AR may directly regulate Nanog transcription [52]. At the post-transcriptional level, we have reported that microRNA-128 may target *Nanog/NanogP8* mRNAs [69]. At the protein level, the ubiquitin ligase SPOP may mediate Nanog degradation via the proteasomal pathway [56, 79, 80] and ERK1-mediated Nanog phosphorylation may also regulate its stability [81]. Recently, the mitotic kinase PLK1 (polo-like kinase 1) was reported to phosphorylate Nanog at Ser41 leading to enhanced stability of Nanog protein, which then drove PCa cell lineage plasticity and anti-androgen resistance via transcriptional and metabolic reprogramming [82]. Interestingly, the amino acid substitutions we observed in NanogP8 mutants in HPCa samples may alter their interactions with these regulatory proteins. This possibility warrants further investigation to fully elucidate the role of Nanog isoforms in PCa development and progression.

As our early [41–43, 67, 68, 72] and current studies have focused on NanogP8 with respect to its tumor-promoting functions in prostate and other cancers, data regarding the expression and functions of other processed Nanog pseudogenes remain limited. Among the *Nanog1* gene and its 11 pseudogenes, only Nanog1, Nanog2 and NanogP8 are free of structural defects and capable of producing full-length Nanog proteins [17–19, 41, 70] (**Figure 1B**). Nanog2, a tandem duplicate of Nanog1, has been shown to be expressed in human leukemia and pluripotent cells [16, 17, 83]. However, like the ES cell-originated *Nanog1*, *Nanog2* transcripts were not detected in HPCa specimens and cancer cell lines we analyzed (**Table 1**), suggesting that *Nanog2* may not be expressed in somatic solid tumors. On the other hand, HPCa specimens highly expressed *Nanog P6* and *NanogP7* transcripts in addition to *NanogP8*. Although all *NanogP6* and *NanogP7* transcripts detected in HPCa specimens harbor premature stop codons and are not predicted to encode a functional Nanog protein, whether these pseudogene transcripts themselves may play a role in PCa remains to be determined. Recent studies have shown that some transcribed pseudogenes function as long non-coding RNAs (lncRNAs), regulating gene expression via epigenetic or post-transcriptional mechanisms [84–86]. Notably, expressed pseudogene transcripts can act as molecular sponges or decoys for microRNAs (miRNAs), due to their similar 3’-untranslated regions (3’-UTRs) with their parent genes. Accumulating evidence suggests that *Nanog1* and *NanogP8* mRNA expression is regulated by miRNAs such as miR-34a, miR-126, miR-149, and miR-128 [69, 70, 74], which downregulate Nanog expression by directly targeting its mRNA. Given that the HPCa-abundant Nanog pseudogene transcripts from *NanogP6* and *NanogP7* retain similar 3’-UTRs to those in *Nanog1* and *NanogP8*, they are likely to share the above-mentioned miRNA-binding sites. Thus, these pseudogene transcripts may function as competing endogenous RNAs or ceRNAs [87], sequestering the ‘suppressive’ miRNAs and thereby relieving repression on *Nanog1/NanogP8* transcription, ultimately promoting Nanog protein expression. In this way, Nanog pseudogene transcripts may cooperate with NanogP8 to promote prostate tumorigenesis.

In summary, here we conducted an in-depth analysis of *Nanog* transcript isoforms in 15 HPCa clinical specimens and various cancer cell lines. The sequencing results indicate that PCa patient specimens do not express *Nanog1* and *Nanog2* but abundantly express protein-coding retrogene *NanogP8* and non-protein coding pseudogene transcripts *NanogP6* and *NanogP7*. Significantly, we demonstrate multiple *NanogP8* isoforms in HPCa patient tumors that manifest differential clonal, sphere and tumor promoting effects in PCa cells (see Graphical Abstract). At present, it remains unclear whether the specific *NanogP8* isoforms (or *NanogP8* SNPs) correlate with tumor grade, stage, castration resistance or metastatic potential. Future studies involving larger patient cohorts will be required to address these important clinically-relevant questions.

## 4. Methods

### 4.1. Cells and reagents

All PCa cell lines (PC3, Du145, PPC-1 and LNCaP) were originally purchased from ATCC (American Type Culture Collection, Manassas, VA) and cultured in the recommended media containing 10% of heat-inactivated FBS (fetal bovine serum). Teratocarcinoma cells (NTERA-2) were purchased from ATCC and cultured in mTeSR^TM^1 medium (STEMCELL Technologies). Nuclear extraction kit was bought from Thermo Scientific. Five anti-Nanog primary antibodies (Abs) were obtained from commercial companies (**Figure 4B**). Secondary and control Abs were purchased from Santa Cruz Biotechnology and ECL reagents were from PerkinElmer, Inc.

### 4.2. Primary human PCa (HPCa) specimens and detection of Nanog transcripts

All HPCa samples were obtained at radical prostatectomy with patients’ consent in accordance with federal and institutional guidelines, which were approved by MD Anderson Cancer Center Institutional Review Board (IRB) (Protocol number LAB04-0498). Tumor pieces, designated as adenocarcinoma by pathological examination and classified as Gleason 6–9, were frozen in liquid nitrogen. The total RNA was isolated from tumor pieces by RNeasy mini kit (Qiagen) and the residual genomic DNA was depleted by on-column digestion with RNase-free DNase. 3 μg RNA was used to generate cDNA with Superscript II reverse transcription (Invitrogen) following the manufacturer’s protocol. The full-length *Nanog* transcript was amplified by using high-fidelity Plus PCR system with the following primers: Forward: 5’-ATGAGTGTGGATCCAGCTTGT-3’ and Reverse: 5’-TCACACGTCTTCAGGTTGCAT-3’.

Control reactions without reverse transcriptase were carried out to exclude DNA contaminations. The PCR products were subcloned into pGEM-T vectors (Promega), and 24 subclones of each HPCa sample were sequenced. The mutations (or SNPs) of *Nanog* transcripts were detected by performing the BLAST analysis between the transcript sequences and the sequences of Nanog pseudogenes in Genbank.

### 4.3. AlwN1 digestion experiments

Total RNA from NTERA-2 or cultured PCa cells (LNCaP, Du145, PC3, and PPC-1), PCa xenografts (LAPC4, and LAPC9), and primary HPCa-derived cells (HPCa157T), as well as 3 lung cancer cell lines (A549, H460 and H1299) were extracted using Qiagen RNasy Plus mini kit (Qiagen). The RT reaction was carried out using SuperScript III first-strand synthesis kit (Invitrogen). PCR were carried out in 25 μl reaction using the following primers: Forward 5’-TTGATACTTTTCCTTCTGGAGGTCC-3’ and Reverse 5’-TCCTTTTTTGCGACACTATTCTCTG-3’. 20 μl of PCR products was digested with AlwN1 according to the manufacturer’s protocol (NEB). Digested PCR products, along with the undigested samples, were analyzed by electrophoresis on 3% agarose gel.

### 4.4. Western blotting (WB)

The whole cell lysates from cells were prepared in RIPA buffer (150 mM NaCl, 50 mM Tris-HCl, pH 7.5, 10 mM EDTA, 1% Nonidet P-40, 0.5% sodium deoxycholate, 0.5% Triton X-100) containing protease inhibitor cocktail, and the protein concentrations were measured via MicroBCA kit (Pierce). 50∼80 µg proteins were analyzed on 4-12.5% gradient SDS-PAGE and gels were transferred to the immobilon-P transfer membrane (Millipore). The membrane was blocked with 5% non-fat milk in TBST (10 mM Tris-HCl, 150 mM NaCl and 0.1% Tween-20) for 1 h at RT and then incubated with primary Abs overnight at 4°C. Membranes were washed three times with TBST buffer, then incubated for 1 h with respective secondary Abs, and developed with ECL Plus WB detection reagent (PerkinElmer).

### 4.5. Site-specific mutagenesis

Using the ‘wild-type’ NanogP8 (NanogP8^WT^) cDNA/pCR2.1 [68] as template, we generated NanogP8^N130S^, NanogP8^L61P^, and NanogP8^D64Y^ mutants (or variants) using the QuickChange II Site-Directed Mutagenesis Kit (Stratagene). After confirmation by sequencing, the BamHI/SalI fragments containing the mutated *Nanog* coding sequences were subcloned into the same sites in pET-28b(+) bacterial expression vector to generate His-tagged fusion protein. Similarly, using NanogP8^N130S^ cDNA/pCR2.1, NanogP8^L61P^ cDNA/pCR2.1 and NanogP8^D64Y^ cDNA/pCR2.1 as DNA templates, we performed the reverse mutagenesis experiments to change these NanogP8 variant proteins back to the NanogP8^WT^ protein by mutating S130N, C182T or T190G. The primers used for the site-directed mutagenesis were listed in **Table S2**.

### 4.6. Bacterial expression and purification of Nanog proteins

cDNA of Nanog1 or NanogP8 variants NanogP8^N130S^, NanogP8^L61P^, and NanogP8^D64Y^ were amplified by PCR using LDF1/LDR1 primers (LDF1 5’-TCTTCCTCTATACTAACATGAGT-3’; LDR1 5-AGGATTCAGCCAGTGTCCA-3’) and cloned in pCR2.1. The EcoRI/NotI or BamHI/SalI fragments containing the *Nanog* coding sequence were then subcloned into the same sites in either pET-28a (+) or pET-28b(+) bacterial expression vector to generate His-tagged fusion proteins. After confirmation of clones by restriction enzyme digestion and sequencing analysis, the plasmids were used to transform BL21 (DE) 3 competent bacterial cells to express the fusion proteins. Nanog protein was induced by 0.5 mM IPTG for 3-6 h at 37°C and bacterial pellet was kept at -80°C.

To purify Nanog proteins, bacterial pellets containing His-tagged Nanog1 or NanogP8 variant proteins were lysed in 8 ml of lysis buffer (50 mM NaH2P04, pH 8.0, 300 mM NaCl, 10 mM Imidazole, protease inhibitor cocktail, and 1 mg/ml lysozyme), sonicated for 10 sec (one pulse), and His-tagged Nanog in the supernatant was purified by nickel beads per manufacturer’s instructions (Qiagen).

### 4.7. Overexpression of NanogP8 isoforms by lentivirus infection

Using pCR2.1 harboring Nanog1, NanogP8^N130S^, NanogP8^L61P^, or NanogP8^D64Y^ cDNA as the DNA template, the full-length Nanog1/NanogP8 cDNAs were amplified with the following primers: Forward primer: 5’-TCGAATTCATGAGTGTGGATCCAGCTTG-3’ (EcoRI) and Reverse primer: 5’-ATCTAGATCACACGTCTTCAGGTTGC-3’ (XbaI). The PCR products were subcloned into pGEM-T vectors and several recombinants for each NanogP8 isoform were chosen for sequencing verification. The confirmed recombinants were double digested by EcoRI and XbaI restriction enzymes. The resulting full-length NanogP8 isoform cDNAs were then subcloned into pLVX-IRES-ZsGreen1 vector (Clontech), which was digested by the same restriction enzymes to generate lentiviral Nanog1 or NanogP8 expression constructs (i.e., pLVX-NanogP8^WT^, pLVX-NanogP8^N130S^, pLVX-NanogP8^L61P^, pLVX-NanogP8^D64Y^ and pLVX-Nanog1). The NanogP8 isoform lentiviruses were produced in HEK293T packaging cells (Clontech) by using Lenti-X HTX packaging system (Clontech, Cat #631251) and the lentivirus titers were determined by counting the numbers of GFP-positive cells with FACS. LNCaP and Du145 cells were infected with the Nanog1/NanogP8 lentiviruses at a multiplicity of infection (MOI) of 20 for 48 hours and then harvested for Western Blotting and immunofluorescence staining.

### 4.8. Clonal, clonogenic and sphere-formation assays, and xenograft tumor experiments

The detailed procedures have been described previously [41,42,68,69]. For clonal assays, cells were seeded at low density (100-300 cells/well) in 6-well plates and allowed to grow until visible colonies were observed. The colonies were counted within 2 weeks. For clonogenic assays, 500-1,000 cells were mixed with Matrigel at 1:1 ratio and then seeded in 6-well plates, and the colonies were enumerated in 2 weeks and photographed. For subcutaneous tumor experiments, 50,000 cells of cultured Du145 infected with lentivirus of NanogP8 species (namely, NanogP8^WT^, NanogP8^N130S^, NanogP8^L61P^, NanogP8^D64Y^) and Nanog1 were mixed with Matrigel at 1:1 ratio and injected subcutaneously (s.c) into the flanks of male NOD/SCID mice. Ten weeks later, mice were sacrificed and tumors were harvested and various parameters were recorded including tumor images, tumor weight, incidence, latency.

## Supporting information

Supplementary Materials Index

## ACKNOWLEDGEMENTS

Work in Tang lab was supported, in part, by grants from the U.S. National Institutes of Health (NIH) National Cancer Institute (NCI) R01CA240290 and 2R01CA240290-06A1, grants from the U.S. Department of Defense (DOD) PC220137 and PC220273, Roswell Park Comprehensive Cancer Center and the NCI Center grant P30CA016056, Roswell Park Alliance Foundation (RPAF) and the George Decker Endowment. We thank other Tang lab members for helpful discussions and suggestions. We apologize to the colleagues whose work was not cited due to space constraint.

## Competing interests

The authors declare no competing interests.

## AUTHOR CONTRIBUTIONS

**B. Liu** and **D.G Tang** led the conceptualization, experimental design and execution, data analysis & interpretation, figures preparation and manuscript writing. **B. Liu** was the primary driver and executer of the project. **J. Shen, Q. Li, X. Chen, K. Rycaj, J. Xiong, R. Liu, R. Mehmood,** and **C. Jeter** assisted in HPCa sample acquisition and in carrying out some experiments. **B. Liu** provided the manuscript draft. **R. Kumar** and **X. Liu** participated in project discussions, data analysis and re-analysis, and manuscript writing. **D.G. Tang** was responsible for the overall development, execution and accomplishment of the project, including project concept, experimental strategies, data acquisition, interpretation and presentation, funding acquisition and manuscript finalization. **D.G. Tang** serves as the lead corresponding author. All authors have read and approved the article.

## ADDITIONAL INFORMATION

**Supplementary information** accompanies the manuscript on *Cancer Heterogeneity and Plasticity* website https://pivotscipub.com/chp

## Notes

### Competing Interest Statement

The authors have declared no competing interest.

