## Supplementary Materials Index for "Multiple isoforms of cancer stemness regulator NanogP8 in clinical prostate cancer specimens and their unique biochemical and oncogenic properties"

Liu B, Kumar R., et al

### Supplementary Materials:

The following supplementary materials are available on the website of this paper:

**Figure S1. All *NanogP7* species expressed in HPCa and cultured PCa cells have a premature stop codon.**

**Figure S2. Reactivity of NanogP8 mutants to 5 anti-Nanog Abs.**

**Figure S3. NanogP8$^{WT}$ and NanogP8 mutants are localized in the nucleus in PCa cells.**

**1. Table S1. Summary of mutations in NanogP8 in various HPCa samples**
**2. Table S2. Primers for site-specific mutagenesis**
