## Supplementary material for "Multiple isoforms of cancer stemness regulator NanogP8 in clinical prostate cancer specimens and their unique biochemical and oncogenic properties": Figure S1

**A**

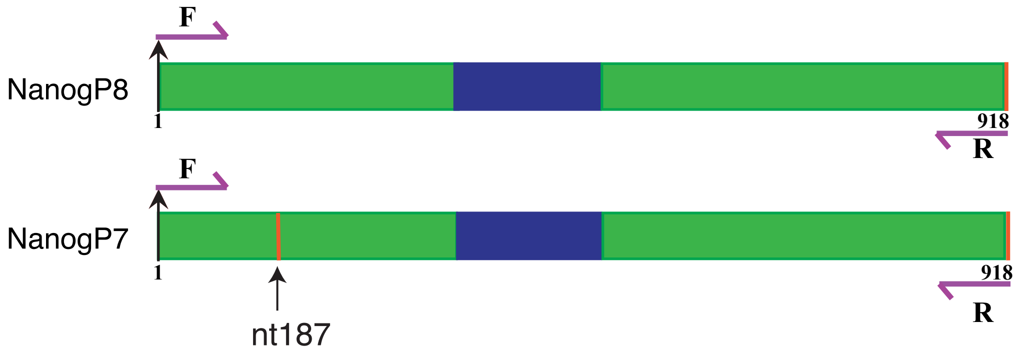

**B**

| | |
|---|---|
| NanogP8 | 171 - GTGGATCTGCTTATTCAGGACAGCCCTGATTCTTCCAC - 209 |
| NanogP7 | 171 - GTGGATCTGCTTATTTAGGACAGCCCTGATTCTTCCAC - 209 |
| HPCa157 | 171 - GTGGATCTGCTTATTTAGGACAGCCCTGATTCTTCCAC - 209 |
| HPCa159 | 171 - GTGGATCTGCTTATTTAGGACAGCCCTGATTCTTCCAC - 209 |
| HPCa161 | 171 - GTGGATCTGCTTATTTAGGACAGCCCTGATTCTTCCAC - 209 |
| HPCa162 | 171 - GTGGATCTGCTTATTTAGGACAGCCCTGATTCTTCCAC - 209 |
| HPCa163 | 171 - GTGGATCTGCTTATTTAGGACAGCCCTGATTCTTCCAC - 209 |
| HPCa165 | 171 - GTGGATCTGCTTATTTAGGACAGCCCTGATTCTTCCAC - 209 |
| HPCa227 | 171 - GTGGATCTGCTTATTTAGGACAGCCCTGATTCTTCCAC - 209 |
| HPCa228 | 171 - GTGGATCTGCTTATTTAGGACAGCCCTGATTCTTCCAC - 209 |
| HPCa231 | 171 - GTGGATCTGCTTATTTAGGACAGCCCTGATTCTTCCAC - 209 |
| DU145 | 171 - GTGGATCTGCTTATTTAGGACAGCCCTGATTCTTCCAC - 209 |
| PC3 | 171 - GTGGATCTGCTTATTTAGGACAGCCCTGATTCTTCCAC - 209 |

Liu & Kumar et al., Figure S1
