## Supplementary figures and images for "Multiple isoforms of cancer stemness regulator NanogP8 in clinical prostate cancer specimens and their unique biochemical and oncogenic properties"

### Figure S2

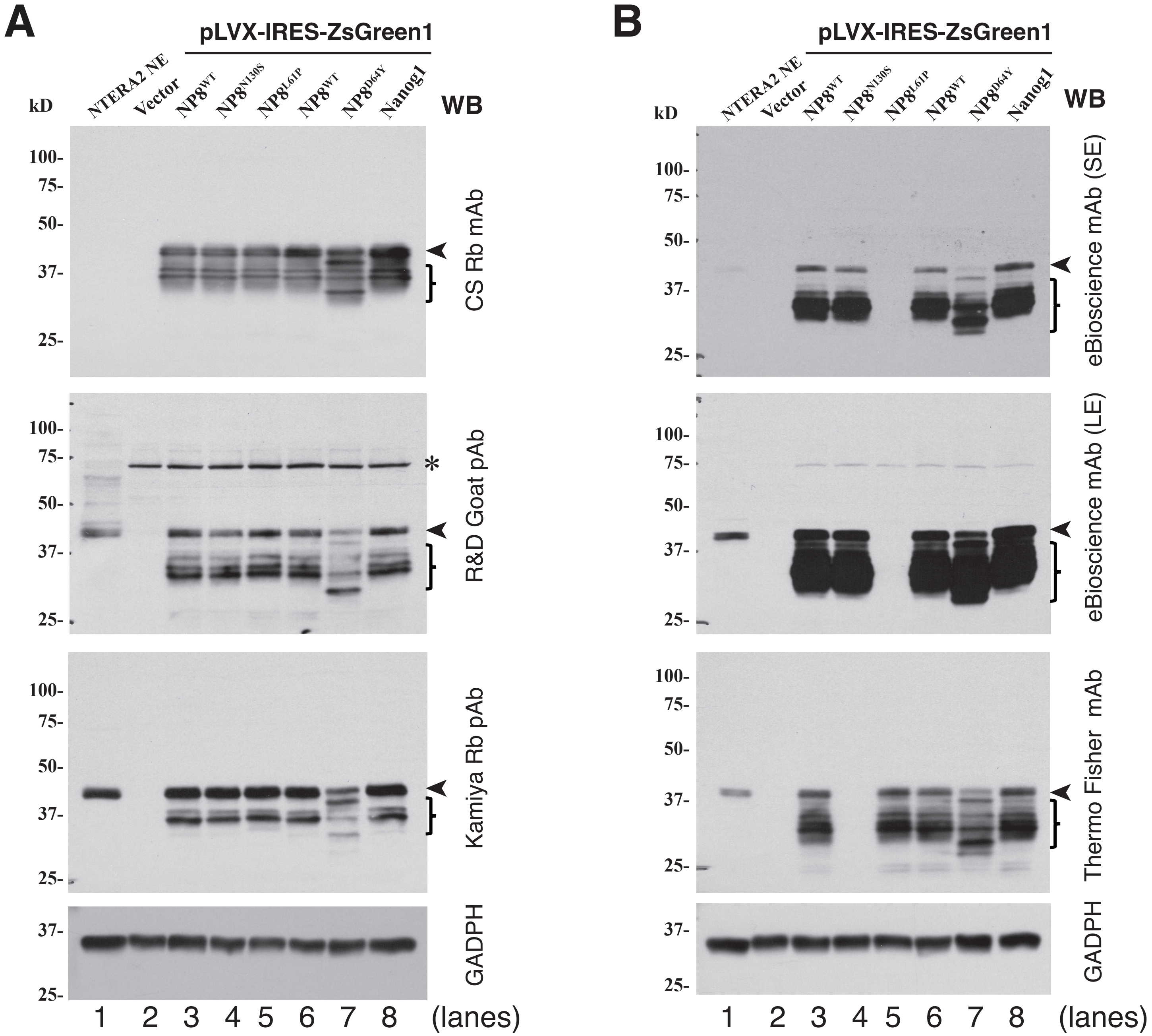


Liu & Kumar et al., Figure S2

### Figure S3

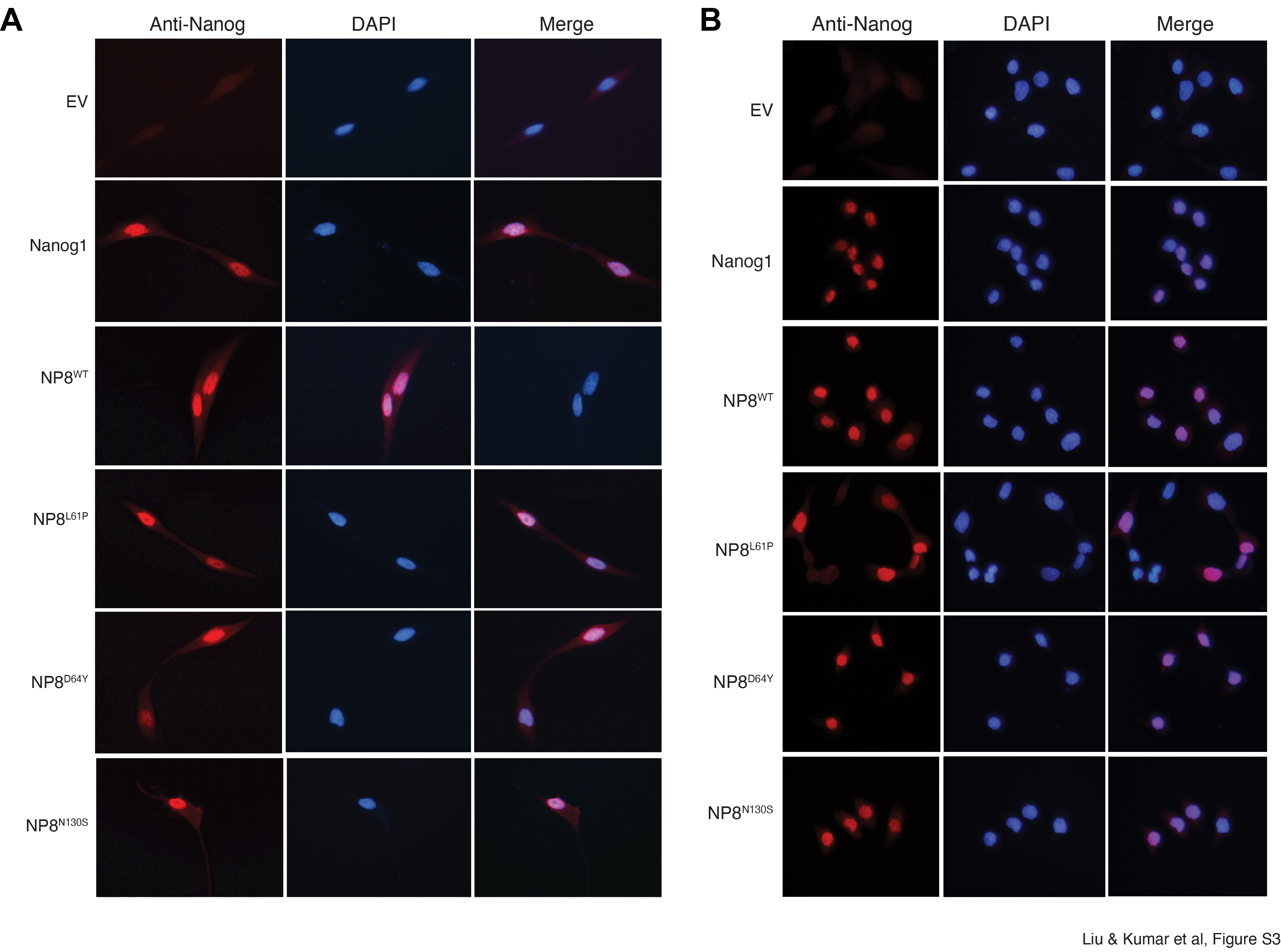
