## Supplementary material for "Multiple isoforms of cancer stemness regulator NanogP8 in clinical prostate cancer specimens and their unique biochemical and oncogenic properties": Table S1

**Table S1: Summary of mutations in NanogP8 in various HPCa samples.**

| Sample / nt position & aa change | HPCa 157 | HPCa 159 | HPCa 161 | HPCa 162 | HPCa 163 | HPCa 165 | HPCa 166 | HPCa 178 | HPCa 216 | HPCa 217 | HPCa 218 | HPCa 219 | HPCa 227 | HPCa 228 | HPCa 231 | Du145 | H1299 | PC3 |
|---|---|---|---|---|---|---|---|---|---|---|---|---|---|---|---|---|---|---|
| 47 C-A<br>GCA-GAA<br>Ala - Glu | | | | | | 1/3 | | 2/4 | 3/7 | 8/15 | | | | 19/23 | | 13/22 | | 2/4 |
| 51 C-T<br>TCC-TCT<br>Ser - Ser | | | | | | | | | | | | | | | | 1/22 | | |
| 63 A-C<br>GAA-GAC<br>Glu - Asp | | | | | | | | | 1/7 | | | | | | 1/19 | | | |
| 71 C-T<br>CCT-CTT<br>Pro - Leu | | | | | | | | 1/4 | 1/7 | | | | | | 1/19 | | | |
| 73 A-G<br>ATG-GTG<br>Met - Val | | | | | | | 1/7 | | | | | | | | | | | |
| 74 T-C<br>ATG-ACG<br>Met- Thr | | | | | | | | | 1/7 | | | | | | | | | |
| 75 G-A<br>ATG-ATA<br>Met - IIe | | | | | | | | | 1/7 | | | | | | | | | |
| 76 C-A<br>CCT-ACT<br>Pro - Thr | 1/13 | | | | | | | | | | | | | | | | | |
| 80 T-C<br>GTG-GCG<br>Val - Ala | 1/13 | | | | | | | | | | | | | | | | | |
| 93 T-C<br>CCT-CCC<br>Pro - Pro | | | | | | | | | | | | | | 1/23 | | | | |
| 101 A-G<br>AAC-AGC<br>Asn - Ser | | | | | | | | | | | 1/12 | | | | | | | |
| 102 C-T<br>AAC-AAT<br>Asn - Asn | 1/13 | | | | | | | | | | | | | | | | | |
| 112 T-C<br>TTG-CTG<br>Leu - Leu | | | | | | | | | | | 1/12 | | | | | | | |
| 116 A-G<br>CAA-CGA | | | | | | | | | | | | | | | | 1/22 | | |

| | | | | | | | | | | | | | | | | | | |
|---|---|---|---|---|---|---|---|---|---|---|---|---|---|---|---|---|---|---|
| Gln - Arg | | | | | | | | | | | | | | | | | | |
| 126 T-C<br>TCT-TCC<br>Ser - Ser | 1/13 | | | | | | | | | | | | | | | | | |
| 133 A-G<br>ATG-GTG<br>Met-Val | | | | | | | | | 1/7 | | | | | | | | | |
| 140 A-G<br>His-Arg | | | | | | | | | | 1/15 | | | | | | | | |
| 142 A-G<br>ACG-GCG<br>Thr - Ala | | | | | | | | | | 1/15 | | | | | | | | |
| 143 C-T<br>ACG-ATG<br>Thr -Met | | | | | | | | | 1/7 | | | | | | | | | |
| 144 G-A<br>ACG-ACA<br>Thr-Thr | 13/13 | 3/3 | 12/12 | 16/16 | 6/6 | 3/3 | 7/7 | 4/4 | 6/7 | 15/15 | 12/12 | 12/12 | 11/11 | 22/23 | 19/19 | 22/22 | 12/12 | 4/4 |
| 145 G-A<br>GAG-AAG<br>Glu - Lys | | | 1/12 | | | | | | | | | | | | | | | |
| 156 T-C<br>TCT-TCC<br>Ser - Ser | | | | | | | | | | | | | 1/11 | | | | 1/12 | |
| 165 T-C<br>CCT-CCC<br>Pro-Pro | | | 1/12 | 2/16 | | | | | | | | | | | | | | |
| 169 T-C<br>TCC-CCC<br>Ser - Pro | | | | | | | | | | | 1/12 | | | | | | | |
| 173 T-C<br>ATG-ACG<br>Met-Thr | | | | 1/16 | | | | | | | | | | | | | | |
| 175 G-A<br>GAT-AAT<br>Asp-Asn | | | | | | | | | | | | | 1/11 | | | | | |
| 182 T-C<br>CTT-CCT<br>Leu - Pro | 2/13 | | | 1/16 | | | | | | | | | | | | | | |
| 187 C-T<br>CAG-TAG<br>Gln-Stop | | | | | | | | | 1/7 | | | | | | | | | |
| 190 G-T<br>GAC-TAC<br>Asp - Tyr | 7/13 | | 10/12 | 8/16 | 6/6 | | | | 1/7 | | | | | | 7/19 | | 12/12 | 2/4 |
| 197 C-A<br>CCT-CAT | 4/13 | | | | | | | | | | | | | | | 1/22 | | |

| | | | | | | | | | | | | | | | | | | |
|---|---|---|---|---|---|---|---|---|---|---|---|---|---|---|---|---|---|---|
| Pro - His | | | | | | | | | | | | | | | | | | |
| 198 T-C<br>CCT-CCC<br>Pro - Pro | | | | | | | | | | | 1/12 | | | | | | | |
| 206 C-T<br>TCC-TTC<br>Ser - Phe | 1/13 | | | | | | | | | | | | | | | | | |
| 208 A-G<br>ACC-GCC<br>Thr - Ala | | | | | | | | | | 1/12 | | | | | | | | |
| 211 A-G<br>AGT-GGT<br>Ser-Gly | | | | | | | | 1/7 | | | 1/12 | | | 1/19 | | | | |
| 212 G-C<br>AGT-ACT<br>Ser - Thr | | | | | | | | 1/7 | | | | | | 1/19 | | | | |
| 221 G-A<br>GGC-GAC<br>Gly - Asp | | | | | | | | 1/7 | | | | | | | | | | |
| 223 A-G<br>AAA-GAA<br>Lys - Glu | | | | | | | | | | | | | | | | 1/12 | | |
| 225 A-G<br>AAA-AAG<br>Lys - Lys | | | | | | | | | | | 1/12 | | | | | | | |
| 246 G-T<br>AAG-AAT<br>Lys - Asn | 13/13 | 3/3 | 11/12 | 16/16 | 6/6 | 3/3 | 7/7 | 4/4 | 7/7 | 15/15 | 12/12 | 12/12 | 11/11 | 23/23 | 19/19 | 22/22 | 12/12 | 4/4 |
| 247 A-G<br>AGT-GGT<br>Ser - Gly | | | | | | | | | | 1/15 | | | | | | | | |
| 251 T-C<br>GTC-GCC<br>Val - Ala | | | | | | | | | | | | | | | 1/19 | | | |
| 260 A-G<br>AAG-AGG<br>Lys - Arg | | | 1/12 | | | | | | | | | | | | 1/19 | | | |
| 261 G-A<br>AAG-AAA<br>Lys - Lys | | | | | | | | | | 2/15 | | | | 1/23 | | | | |
| 262 G-A<br>GAA-AAA<br>Glu - Lys | | | | | | | | | | | | | | | | 1/22 | | |
| 266 A-G<br>GAC-GGC<br>Asp - Gly | | | | | | | | | | | 1/12 | | | | | | | |
| 267 C-T<br>GAC-GAT | | | | | | | | | | | | | | | | | 1/12 | |

| | | | | | | | | | | | | | | | | | |
|---|---|---|---|---|---|---|---|---|---|---|---|---|---|---|---|---|---|
| Asp - Asp | | | | | | | | | | | | | | | | | |
| 270 G-A<br>AAG-AAA<br>Lys - Lys | | | | | | | | | 1/7 | | | | | | | | |
| 275 C-T<br>CCG-CTG<br>Pro - Leu | | | | | | | | | | | 1/12 | | | | | | |
| 276 G-A<br>CCG-CCA<br>Pro-Pro | | | 1/12 | 2/16 | | | | | | | | | | | | | |
| 289 A-G<br>AAG-GAG<br>Lys - Glu | | | | 1/16 | | | | | | | | | | | | | |
| 290 A-G<br>AAG-AGG<br>Lys - Arg | | | 1/12 | | | | | | | | | | | | | | |
| 298 A-G<br>ACT-GCT<br>Thr - Ala | | | | | | | | | | | | | | | | 1/22 | |
| 300 T-A<br>ACT-ACA<br>Thr - Thr | | | | | | | | | | | | | | | | 1/22 | |
| 302 T-C<br>GTG-GCG<br>Val - Ala | | | | | | | | 1/4 | | | | | | | | | |
| 303 G-A<br>GTG-GTA<br>Val - Val | | | | 1/16 | | | | | | | | | | | | | |
| 309 T-C<br>TCT-TCC<br>Ser - Ser | | | | | | | | | 1/7 | | | | | | | | |
| 313 A-G<br>ACC-GCC<br>Thr - Ala | | | | | | | | | 1/7 | | | | | | | | |
| 314 C-T<br>ACC-ATC<br>Thr - IIe | | | | | | | | | | | 1/12 | | | | | | |
| 320 T-C<br>CTG-CCG<br>Leu - Pro | | | | | | | | | | | | 1/12 | | | | | |
| 324 T-C<br>TGT-TGC<br>Cys - Cys | | | | | | | | | | | 1/12 | | | | | | |
| 327 A-G<br>GTA-GTG<br>Val - Val | | | | | | | | | | | | | | | | | 1/12 |
| 333 T-C<br>AAT-AAC | | | | 1/16 | | | | | | | | | | | | | |

| | | | | | | | | | | | | | | | | | |
|---|---|---|---|---|---|---|---|---|---|---|---|---|---|---|---|---|---|
| Asn - Asn | | | | | | | | | | | | | | | | | |
| 339 A-G<br>AGA-AGG<br>Arg - Arg | | | | 1/16 | | | | | | | | | | | | | |
| 346 A-G<br>AGA-GGA<br>Arg -Gly | | | | | | | | | | | | | | | 1/19 | | |
| 356 A-G<br>TAC-TGC<br>Tyr - Cys | | | | | | | | | | | 2/12 | | 1/11 | | | | 1/12 |
| 377 A-G<br>CAA-CGA<br>Gln - Arg | | | | | | | | | | 1/15 | | | | | | 1/22 | |
| 387 C-T<br>TCC-TCT<br>Ser - Ser | | | | | | | | | | 1/15 | | | | | | | |
| 395 T-C<br>CTG-CCG<br>Leu - Pro | | | | | | | | | | | | 1/12 | | | | | |
| 399 C-T<br>AAC-AAT<br>Asn - Asn | | | | | | | | | | | | | | | | 1/22 | |
| 407 A-G<br>TAC-TGC<br>Tyr-Cys | | | | | 1/6 | | | | | | | | | | | | |
| 416 T-C<br>GTG-GCG<br>Val - Ala | | | | | | | | | | | | | 1/11 | | | | |
| 421 A-G<br>ACC-GCC<br>Thr - Ala | | | | | | | | | | 1/15 | | | | | | | |
| 422 C-T<br>ACC-ATC<br>Thr - IIe | | | | | | | | | | | | 2/12 | | | | | |
| 428 T-C<br>TTC-TCC<br>Phe - Ser | | | | | | | | | | | 1/12 | | | | | | |
| 437 A-G<br>CAG-CGG<br>Gln - Arg | | | | | | | | | | 1/15 | | | | | | | |
| 442 A-G<br>ATG-GTG<br>Met - Val | | | 1/12 | | | | | | | | | | | | | | |
| 447 A-G<br>AAA-AAG<br>Lys - Lys | | | | | | | | | | | 1/12 | | | | | 1/22 | |
| 452 A-G<br>AAG-AGG | | | | | | | | | | | | | | | | | |

| | | | | | | | | | | | | | | | | | | |
|---|---|---|---|---|---|---|---|---|---|---|---|---|---|---|---|---|---|---|
| Lys - Arg | | | | | | | | | | | | | | | | | | |
| 454 A-G<br>AGG-GGG<br>Arg - Gly | 1/13 | | | | | | | | | | | | | | | | | |
| 455 G-A<br>AGG-AAG<br>Arg - Lys | | | | | | | | | | | 1/12 | | | | | | | |
| 458 G-A<br>TGG-TAG<br>Trp - Stop | 1/13 | | | | | | | | | | | | | | | | | |
| 460 C-T<br>CAG-TAG<br>Gln - Stop | | | | | | | | | 1/7 | | 1/12 | | | | | | | |
| 463 A-G<br>AAA-GAA<br>Lys - Glu | | | 1/12 | | | | | | | | | | | | | | | |
| 465 A-G<br>AAA-AAG<br>Lys - Lys | | | | | | | | | | | 1/12 | | | | | | | |
| 474 G-A<br>TGG-TGA<br>Trp - Stop | | | | | | | | | | | | | | | | | | |
| 477 G-A<br>CCG-CCA<br>Pro - Pro | | | 1/12 | | | | | | | | | | | | | | | |
| 479 A-G<br>AAG-AGG<br>Lys - Arg | | | | | | | | | | | | | | | | | 1/12 | |
| 488 A-G<br>AAT-ATG<br>Asn - Met | | | | | | | | | | | | | | | | 1/22 | | |
| 492 T-C<br>GGT-GGC<br>Gly - Gly | | | | | | | | | | | | | | | | | | |
| 498 G-T<br>ACG-ACT<br>Thr - Thr | | | | | | | | | | | | | | | | | | |
| 507 C-T<br>GCC-GCT<br>Ala - Ala | | | | | | | | | | | | | | | | 1/22 | | |
| 531 T-C<br>CTT-CTC<br>Leu - Leu | 13/13 | 3/3 | 11/12 | 16/16 | 6/6 | 3/3 | 7/7 | 4/4 | 7/7 | 15/15 | 12/12 | 12/12 | 11/11 | 23/23 | 19/19 | 22/22 | 12/12 | 4/4 |
| 534 C-T<br>TAC-TAT<br>Tyr - Tyr | | | | | | | | | | | | | | | | 1/22 | | |
| 535 T-C<br>TCT-CCT | | | | | | | | | | | | | | | | | | 1/4 |

| | | | | | | | | | | | | | | | | | | |
|---|---|---|---|---|---|---|---|---|---|---|---|---|---|---|---|---|---|---|
| Ser - Pro | | | | | | | | | | | | | | | | | | |
| 537 T-C<br>TCT-TCC<br>Ser - Ser | | | | | | 1/3 | 1/7 | | | | | | | | | | | |
| 542 A-G<br>TAC-TAG<br>Tyr - **Stop** | | | | | | | | | | | | | | | | 1/22 | | |
| 547 C-T<br>CAG-TAG<br>Gln - **Stop** | | | 1/12 | | | | | | | | | | | | | | | |
| 548 A-G<br>CAG-CGG<br>Gln - Arg | | | 1/12 | | | | | | | | | | | | | | | |
| 550 G-A<br>GGA-AGA<br>Gly-Arg | | | | | | | | | | | | | | | | 1/22 | | |
| 552 A-G<br>GGA-GGG<br>Gly - Gly | 6/13 | | 8/12 | 11/16 | 6/6 | | | 1/4 | | | | | | | 7/19 | | 12/12 | 1/4 |
| 562 A-G<br>AAC-GAC<br>Asn - Asp | | | | | | | | | | | 1/12 | | | | | | | |
| 565 C-A<br>CCG-ACG<br>Pro - Thr | | | 1/12 | | | | | | | | | | | | | | | |
| 566 C-T<br>CCG-CTG<br>Pro - Leu | | | 1/12 | | | | | | | 1/15 | | | | | | | | |
| 574 A-G<br>AAC-GAC<br>Asn - Asp | | | | | | 1/3 | | | | | | | | | | | | |
| 576 C-T<br>AAC-AAT<br>Asn - Asn | 1/13 | | | 1/16 | | | | | | | | | | | | | | |
| 577 C-T<br>CTT-TTT<br>Leu - Phe | | | | | | | | | | | | | | | 1/19 | | | |
| 589 A-G<br>AGC-GGC<br>Ser - Gly | | | | | | | | | | | | | | | 1/19 | | | |
| 590 G-A<br>AGC-AAC<br>Ser - Asn | | | | | | | | | | 1/15 | | 2/12 | | 1/23 | | | | |
| 598 A-G<br>ACC-GCC<br>Thr-Ala | | | | | | | | | | | | | 1/11 | | | | | |
| 601 T-C<br>TGG-CGG | 1/13 | | | | | | | | | | | | | | | | | |

| | | | | | | | | | | | | | | | | | |
|---|---|---|---|---|---|---|---|---|---|---|---|---|---|---|---|---|---|
| Trp - Arg | | | | | | | | | | | | | | | | | |
| 602 G-A<br>TGG-TAG<br>Trp - **Stop** | | | | 1/16 | | | | | | | | | | | | | |
| 607 A-G<br>AAT-GAT<br>Asn - Asp | | | | | | | | | | 1/15 | | | | | | | |
| 608 A-T<br>AAT-ATT<br>Asn - IIe | | | | | | | | | | 1/15 | | | | | | | |
| 612 A-G<br>TCA-TCG<br>Ser - Ser | | | | | | | | | | | | | | | | | 1/12 |
| 619 A-G<br>AGC-GGC<br>Ser - Gly | | | | | | | | | | | 1/12 | | | | | | |
| 626 A-G<br>CAG-CGG<br>Gln - Arg | | | | | | | | | | | | | | | 1/19 | | 1/12 |
| 629 C-T<br>ACC-ATC<br>Thr - IIe | | | | | | | | | 3/7 | | | | | | | | |
| 630 C-T<br>ACC-ACT<br>Thr - Thr | 1/13 | | | | | | | | | | | | | | | | |
| 638 T-G<br>ATC-AGC<br>IIe - Ser | | | | | | | | | | 1/15 | | | | | | | |
| 639 C-T<br>ATC-ATT<br>IIe - IIe | | | | | | | | | | | | | | | | | 1/12 |
| 646 T-C<br>TGG-CGG<br>Try-Arg | | | | | | | | | | | | | | 1/23 | | 1/22 | |
| 648 G-A<br>TGG-TGA<br>Trp - **Stop** | | | | | | | | | | | 1/12 | | | | | | |
| 650 G-A<br>AGC-AAC<br>Ser - Asn | | | | | | | | | | | | | | | | | 1/12 |
| 653 A-G<br>AAC-AGC<br>Asn - Ser | | | | | | | | | | | | | | | | 1/22 | |
| 655 C-T<br>CAC-TAC<br>His - Tyr | 1/13 | | | | | | | | | | | | | | | | |
| 665 A-G | | | | | | | | | | 1/15 | | | | | | | |

| | | | | | | | | | | | | | | | | | |
|---|---|---|---|---|---|---|---|---|---|---|---|---|---|---|---|---|---|
| AAC-AGC<br>Asn - Ser | | | | | | | | | | | | | | | | | |
| 667 A-G<br>ACT-GCT<br>Thr - Ala | | | | | | | | | 1/7 | | | | | | | | |
| 669 T-C<br>ACT-ACC<br>Thr - Thr | | | | | | | | | | | | 1/12 | | | | | |
| 683 C-T<br>ACC-ATC<br>Thr - IIe | | | | | | | | | 1/7 | | | | | | | | |
| 687 A-G<br>CAA-CAG<br>Glu-Glu | | | | | | | | | | | | | | | 1/19 | | |
| 697 A-G<br>AAT-GAT<br>Asn - Asp | | | | | | | | | | | | 1/12 | | | | | |
| 702 G-C<br>CAG-CAC<br>Gln - His | 2/13 | | | | | | | | | | | | | | | | |
| 712 A-G<br>AGT-GGT<br>Ser - Gly | | | | | | | | | | | | | | | | 1/22 | |
| 714 T-C<br>AGT-AGC<br>Ser - Ser | | | | 1/16 | | | | | | | | | | | | | |
| 725 A-G<br>AAC-AGC<br>Asn - Ser | | | | 1/16 | | | | | | | | | | | | | |
| 728 G-A<br>TGT-TAT<br>Cys - Tyr | | | | | | | | | | | | | | | 1/19 | | |
| 745 C-T<br>CAG-TAG<br>Gln - **Stop** | | | | 1/16 | | | | | | | | | | | | | |
| 746 A-G<br>CAG-CGG<br>Gln - Arg | | | | | | | | | | | | | | | | 1/22 | 2/12 |
| 751 T-C<br>TGC-CGC<br>Cys - Arg | | | 1/12 | | | | | | | | | | | | | | |
| 754 A-C<br>ATG-CTG<br>Met - Leu | | | | | | | | | | | | | | 3/23 | | | |
| 756 G-A<br>ATG-ATA<br>Met-Ile | | | | | | | | | | 1/15 | | | | | | | |

| | | | | | | | | | | | | | | | | | | |
|---|---|---|---|---|---|---|---|---|---|---|---|---|---|---|---|---|---|---|
| 759 G-C<br>CAG-CAC<br>Gln - His | 13/13 | 3/3 | 12/12 | 16/16 | 6/6 | 3/3 | 7/7 | 4/4 | 7/7 | 15/15 | 12/12 | 12/12 | 11/11 | 23/23 | 19/19 | 22/22 | 12/12 | 4/4 |
| 760 T-C<br>TTC-CTC<br>Phe - Leu | | | | | | | | | | | 1/12 | | | | | | | |
| 765 G-A<br>CAG-CAA<br>Gln - Gln | | | | | | | | | | | | 1/12 | | | | | | |
| 791 A-G<br>GAG-GGG<br>Glu - Gly | | | | 1/16 | | | | | | | | | | | | | | |
| 793 G-A<br>GCT-ACT<br>Ala - Thr | 1/13 | | | | | | | | | | | | | | | | | |
| 798 T-C<br>GCC-GCT<br>Ala - Ala | | | | 3/16 | | | | | | | | | | | | | | |
| 799 T-A<br>TTG-ATG<br>Leu - Met | | | | 1/16 | | | | | | | | | | | | | | |
| 806 C-T<br>GCT-GTT<br>Ala - Val | | | | | | | | | | | | 1/12 | | | | | | |
| 809 C-T<br>GCT-GTT<br>Ala - Val | | | | | | | | | | | | | 1/11 | | | | | |
| 846 G-A<br>AGG-AGA<br>Arg - Arg | | | | | | | | | | | | | | | | 1/22 | | |
| 853 A-G<br>AGT-GGT<br>Ser - Gly | | | | | | | | | | | | 1/12 | | | | | | |
| 873 T-C<br>GAT-GAC<br>Asp - Asp | | | | 1/16 | | | | | | | | | | | | | | |

This Table gives a detailed summary of the mutations detected in NanogP8 (NP8) transcripts from the 15 HPCa samples and 3 cancer cell lines (indicated on the top row). In total, 24 subclones from each sample underwent sequencing analysis. The left-most column describes all NP8 mutations (including their location, codon, and predicted amino acid (aa) changes. The fraction in the grid represents the ratio of mutation(s) to the total NP8 transcripts. For example, the number “1/3” in the sixth grid of the second row indicates that 3 transcripts of NP8 were detected in the sample HPCa 165 from total 24 subclones, of which only one NP8 transcript has the mutation (C-A) at position of 47. Du145 and PC3 are prostate cancer cell lines while H1299 is a lung cancer cell line.
