## Supplementary material for "Multiple isoforms of cancer stemness regulator NanogP8 in clinical prostate cancer specimens and their unique biochemical and oncogenic properties": Table S2

### Table S2 Primers for site-specific mutagenesis

| Site-specific mutagenesis | Primers | Amino acid change |
|---|---|---|
| Nanog cDNA nt sequence<br>(HPCa1, nt168 – nt203) | 5'- CTCCATGGATCTGC**T**TATTCAG**G**ACAGCCCTGATTC -3<br>3'-GACCTACCTAGACG**A**ATAAGTC**C**TGTCGGGACTAAG -5' | L61 (Leu61)<br>D64 (Asp64) |
| HPCa1 to HPCa6<br>(T182C) | For 5'-CTCCATGGATCTGC***c***TATTCAGGACAGCCCTGATTC -3'<br>Rev 3'-GACCTACCTAGACG***g***ATAAGTCCTGTCGGGACTAAG -5' | L61P (Leu61Pro) |
| HPCa1 to MCF7<br>(G190T) | For 5'-CTCCATGGATCTGCTTATTCAG***t***ACAGCCCTGATTC -3'<br>Rev 3'-GACCTACCTAGACGAATAAGTC***a***TGTCGGGACTAAG -5' | D64Y (Asp64Tyr) |
| HPCa6 to HPCa1<br>(C182T)<br>MCF7 to HPCa1<br>(T190G) | For 5'-CTCCATGGATCTGC***t***TATTCAG***g***ACAGCCCTGATTC -3'<br>Rev 3'-GACCTACCTAGACG***a***ATAAGTC***c***TGTCGGGACTAAG -5' | P61L (Pro61Leu)<br><br>Y64D (Tyr64Asp) |
